# AD-genes and Aging Increase Count and Size of Lipid Droplets, Accompanied by Accumulation of Neutral Lipids Across Compartments in Hippocampal Neurons

**DOI:** 10.1101/2025.07.01.661999

**Authors:** Balam Benítez-Mata, Joshua McWhirt, Ricardo Santana, Gregory Brewer, Michelle A. Digman

## Abstract

Lipid homeostasis plays a crucial role in neuronal function, yet its dynamics during aging and neurodegenerative diseases remain poorly understood. Our study unveils critical age-related changes in lipid polarity and lipid droplet characteristics in hippocampal neurons from non-transgenic (NTg) mice and from an Alzheimer’s disease-model (3xTg-AD). Using advanced spectral imaging and phasor analysis techniques, we tracked lipid polarity with Nile Red in vitro across various cellular compartments and quantified lipid droplet features. We discovered that NTg neurons exhibit a progressive increase in global lipid polarity from young to middle age, followed by a slight decrease in old age. This pattern suggests that neurons actively regulate their lipid composition throughout the lifespan, potentially in response to changing cellular needs. In contrast, AD-like (or 3xTg-AD) neurons fail to show this age-related increase in lipid polarity, instead displaying a consistent reduction in lipid polarity across all ages. Lipid droplet analysis revealed a transient accumulation of larger droplets in middle-aged NTg neurons, while AD-transgenic neurons showed early and persistent increases in lipid droplet size and number. Principal component analysis uncovered coordinated changes in lipid polarity and droplet characteristics, highlighting distinct patterns of lipid partitioning in NTg and AD-affected neurons. These findings suggest that AD-associated genetic modifications disrupt normal age-related adaptations in lipid metabolism and organization. Our results provide new insights into the complex interplay between lipid homeostasis, aging, and AD-genotypic stress. Understanding these dynamics may open new avenues for developing therapeutic strategies to maintain neuronal health and potentially slow AD progression.

## INTRODUCTION

Approximately 50% of the brain’s dry weight comprises lipids ^1^. Lipids fulfill various roles in the cell to maintain membrane functionality, signaling responses to organelle stress, provide structural integrity, and serve as energy substrates ^2–5^. These roles are typically negatively affected during brain aging, which is characterized by the functional and structural decline of various cells and organs, leading to impairments in processes such as cognition and memory. Many of these alterations are linked to the neuronal membrane lipid profile ^3^. Lipid profiling studies in brain aging have found region-specific changes in some major lipid groups ^6,7^. Some of these changes constitute major risk factors for pathologies including Alzheimer’s disease (AD) ^3,5^.

In AD, alterations in lipid groups primarily cholesterol ^8^, phospholipids, and sphingolipids ^9^ results in disruption of lipid homeostasis, impairing the cell’s ability to maintain lipid membrane composition and functionality. Thus, lipid metabolism becomes a key factor in disease development, which exacerbates lipid alterations observed during NTg brain aging ^6^. Other studies have detailed lipid profiles across different brain regions and cell types aiding in the understanding of early neurodegenerative processes ^2,10^. The hippocampus, being a key memory processing unit in the brain, is of particular interest in understanding cognitive functions and neurological disorders. Given the region’s crucial role in learning, memory consolidation, and its elevated vulnerability during aging and AD ^11,12^. Studying these neurons poses significant challenges that demand specialized methods. The hippocampus’ deep brain location complicates access, requiring advanced techniques. Additionally, neuronal plasticity and context-dependent changes in morphology and lipid metabolism further complicate their analysis.

After lipid synthesis, various lipids are translocated to different cell organelles where their composition correlates with organelle functions ^13–15^. Among cell organelles, lipid droplets stand out as the most lipid-rich, consisting of a hydrophobic core containing neutral lipids such as triglycerides and cholesteryl esters, surrounded by a monolayer of polar lipids, predominantly phospholipids and lipoproteins, which regulate their dynamics ^16–19^. Lipid droplets play a crucial role in neuronal survival, particularly under conditions of aging, oxidative stress, and inflammation, which makes Lipid droplets a dynamic and constantly changing organelle leading to several types of Lipid droplets populations ^19–23^. Conventional microscopy techniques fall short in capturing the structural diversity of heterogeneous cellular populations, as they cannot accurately resolve the sizes and proportions of distinct subgroups. To address this, we used PLICS (Phasor analysis of Local Image Correlation Spectroscopy), a rapid, unbiased, and fit-free method to quantify the size and number of lipid droplets in neuronal cells with high precision ^24^.

To characterize and understand lipid alterations, biochemical analyses, fluorescence imaging using lipid-sensitive probes, and metabolomic studies have been employed to monitor lipid fluidity, polarity, and composition in neuronal cells and brain tissues ^3,6,7,25–29^. While biochemical analyses reveal lipid composition, imaging is essential for mapping subcellular features and detecting lipid changes in physiological states. Nile Red, a lipophilic solvatochromic dye, is widely used for probing lipid environments due to its polarity-sensitive emission—shifting from 580 nm in non-polar to 640 nm in polar environments—allowing global lipid mapping within cells using a single fluorescent probe ^25,30,31^. To monitor changes in living cells, we and others have applied the spectral phasor approach to hyperspectral imaging of environment-sensitive dyes ^32–34^. This rapid, model-free method enables unmixing of complex spectral signatures and spatial mapping of distinct cellular compartments based on their phasor profiles ^35^.

Our findings reveal distinct patterns of lipid polarity and lipid droplet characteristics in normal adult mouse neurons across the age-span versus pathologic progression in those from an AD mouse model. The age-dependent increase in lipid polarity and average number of lipid droplets observed in NTg neurons contrasts sharply with the disrupted lipid polarity and abnormal lipid droplet accumulation in AD-transgenic neurons. Multiplexing the hyperspectral imaging by applying the spectral phasor approach and PLICS in age-related and AD studies provides a powerful tool for unraveling the complex interplay between lipid polarity, lipid droplet dynamics, and neuronal health. These results not only highlight the potential of lipid polarity and lipid droplet related genes as biomarkers for early AD detection but also open new avenues for therapeutic interventions targeting lipid metabolism in neurodegenerative diseases. Future investigations building on these findings may contribute to a more comprehensive understanding of neurological processes, potentially informing new research directions and finding novel therapeutic approaches. Ultimately, this expanded perspective on lipid roles could enhance our knowledge of Alzheimer’s disease and age-related neurological disorders, opening avenues for improved management strategies.

## RESULTS

### Nile red as a sensitive polarity reporter in solutions and cells

In Figure 1A we schematically show nile red’s fluorescence emission spectrum in the presence of three different lipid polarity environments in a lipid bilayer. In neutral lipid environments, comparable to cholesterol and triglycerides fat droplets, the spectrum is centered at 583 ± 46 nm, corresponding to the yellow-gold regions of the light spectrum. In an environment abundant with polar lipids, such as a phospholipid bilayer, the emission predominantly occurs at wavelengths of 637 ± 52 nm, corresponding to the red and far-red regions of the light spectrum. In the presence of a mixture of neutral and polar lipid environments, the emission spectrum shifts to wavelengths in between polar and neutral lipid spectral emissions, ranging from ∼583 to ∼637 nm. Nile red’s sensitivity to environmental polarity stems from its increased dipole moment upon excitation. This enhancement in the dipole moment triggers a time-dependent process called dipolar relaxation. During this process, the dipoles of the surrounding solvent molecules reorient in response to nile red’s altered electronic state. This reorientation of solvent dipoles is what allows nile red to effectively sense the polarity of its environment. (Supplementary Figure 1A) ^36,38,40^. This photophysical effect is characteristic of other solvatochromic dyes including LAURDAN and the DAN probes used to characterize the fluidity of membranes^42^. We specifically chose nile red for our studies due to its exceptional sensitivity to environmental hydrophobicity, allowing us to distinguish between various lipid environments with remarkable precision. Nile red’s ability to easily permeate cell membranes and stain both intracellular lipid droplets and membranes enhances its versatility in our lipid biology studies.

**Figure 1.**
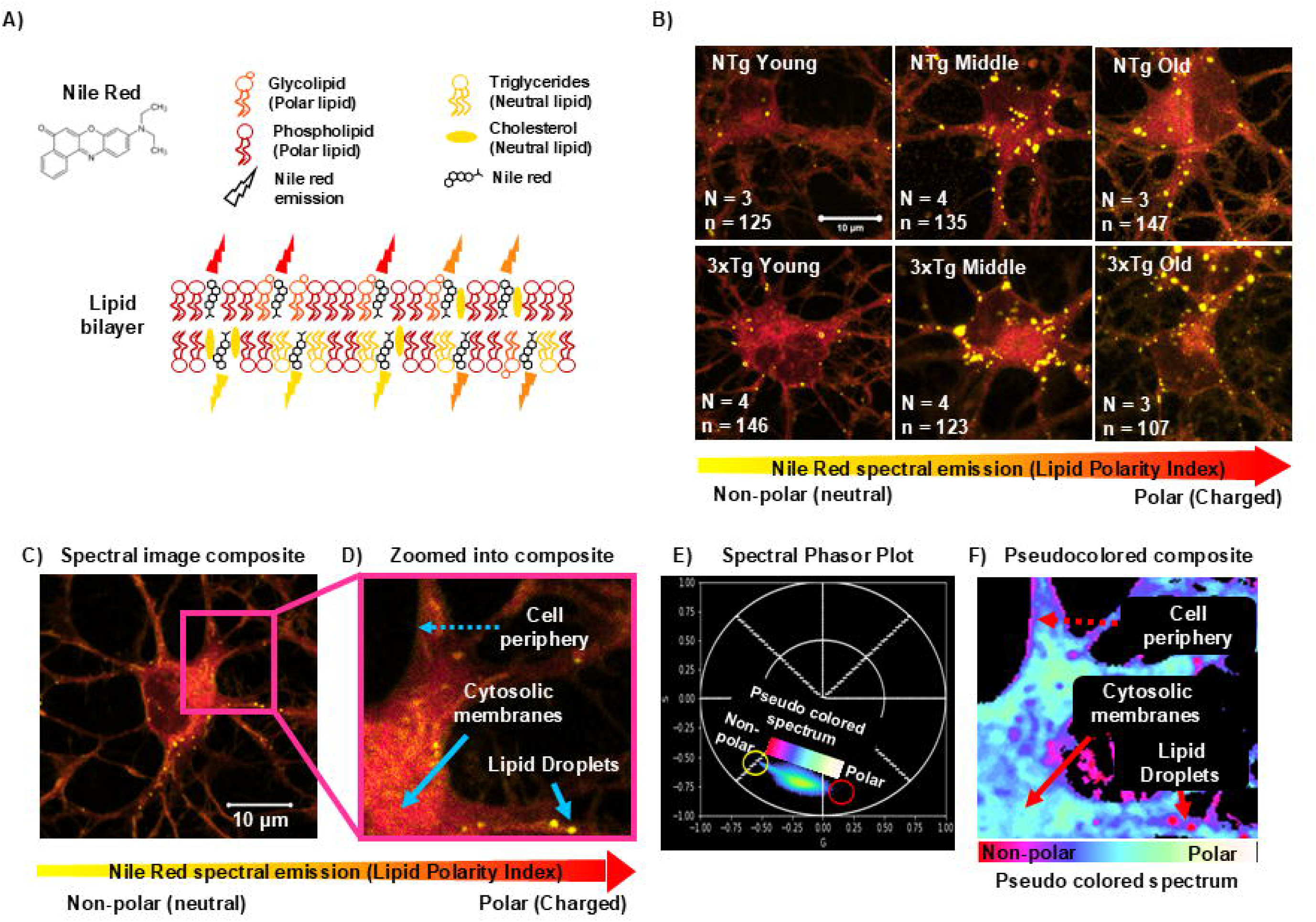
Nile Red photophysics in neuronal lipid environments unveils lipid poplarity gradients in cytosolic and plasma membranes highlight cellular compartment heterogeneity. A) Schematic representation of NR insertion into the lipid bilayer and a simplified photodynamiesfluorescence emission in the presence of different types of lipids. 8) Representative composite spectral images for young, middle and old neurons for both genotypes, NTg and 3xTg, “N” represents animal replicates, while “n” total number of analyzed images. C) Representative enlarged composite image. D) Enlarged composite image. Squared dotted arrow points to a portion of the cell periphery. Solid arrow points to a cytosolic portion of the cell. E) Spectral phaser plot with pseudocolored scale to highlight composite pixels with their respective phaser location. F) Pseudocolored zoomed in ROI of composite image. Squared dotted arrow points to a portion of the cell periphery. Solid arrow points to a cytosolic portion of the cell. Scale bar is 10 micrometers.

We first tested nile red’s ability to sense environmental polarities in organic solvents. We use multiple solvents ranging low to high polarities based on their dielectric constant (hexane ε = 1.88, chloroform ε = 4.81, acetone ε = 20.7, methanol ε = 32.7, and water ε = 78.4). Nile red’s emission displays a red-shift as solvent polarity increases, in Supplementary Figure 1B we show the real-color emission of nile red in each solvent. When the spectral data from each nile red in solution condition is transformed to the phasor space, a clear separation of phasor clouds is evident, showing the ability of nile red to sense different polarity environments and the phasor approach to successfully separate them as a fit-free method to retrieve spectral signatures.

Neurons are one of the most lipid-rich cell types because of their extensive axonal and dendritic membrane specialization^37^. In this work we used nile red to probe the lipid polarity of cellular membranes within neurons, providing valuable information regarding membrane organization, normal cellular composition, and potential alterations associated with aging and neurodegenerative disease. Figure 1B illustrates the range of spectral emissions of nile red in hippocampal neurons of varying ages and genotypes, including non-transgenic (NTg) and triple-transgenic Alzheimer’s disease (3xTg-AD) models. In these neurons, nile red reveals two primary environments: a polar environment with a spectral peak centered at 637 nm (appearing red) observed in the cytosol, and a neutral environment with a spectral peak centered at 583 nm (appearing intense yellow-gold). The latter highlights spherical structures attributed to lipid droplets, consistent with nile red’s previously documented high affinity for lipids, particularly lipid droplets (LDs)^30^ (Supplementary Figure 2). Confirmation of nile red’s yellow-gold emission staining lipid droplets but not other circular-shape organelles such as recycling vesicles or lysosomes is seen by co-staining SH-SY5Y cells with nile red and LipiBlue, a LDs-specific fluorescent probe ^39^(Supplementary Figure 3).

Similarly, when the spectral emission of nile red in neurons is transformed into the spectral phasor space, very non-polar and polar regions within the cell display unique phasor signatures, phasors corresponding to non-polar or neutral environments and polar environments are displayed in opposite directions with respect to each other in the phasor plot (Supplementary Figure 3).

### Lipid Polarity Gradients in Cytosolic and Plasma Membranes Highlight Cellular Compartment Heterogeneity

A full overview of a neuron seems to display only two lipid environments, reveals a gradient of lipid environments within the cytoplasm, ranging from neutral (yellow-gold) to polar (red), resulting in a gradual transition effect in certain regions (Figure 1C). Upon higher magnification of a region of interest (ROI) encompassing both the cell edge and cytoplasm, we observed distinct lipid polarity patterns specific to the cell periphery and cytosolic membranes (Figure 1D). These observations underscore the heterogeneity of lipid environments within different cellular compartments and highlight the importance of spatial resolution in lipid polarity studies.

To better visualize these differences, we applied pseudocoloring based on lipid polarity levels, mapping the neutral-to-polar spectrum using a high-contrast color palette in both the spectral phasor plot and the composite image (Figure 1E & Supplementary Figure 6A). This approach produces a high-contrast “spectral phasor lambda” color coded image where highly neutral regions appear red-magenta (emission peak at 583 nm), and highly polar regions appear yellow-white (emission peak at 637 nm) (Figure 1F & Supplementary Figure 6B).

Polarity frequency histograms, cross-section lines derived from the phasor lambda plots, confirm that the cell periphery (plasma membrane (PM) and cytosolic membranes have distinct polarity values (Supplementary Figure 6C-D). To analyze how each compartment is impacted during aging, we created a cell periphery mask that includes the plasma membrane and part of the inner cytosolic leaflet. We then subtract this mask from the whole-cell mask using logical operations, followed by another subtraction to remove LD. The two resulting binary images are combined to generate a final binary mask that excludes the cell periphery and LDs, effectively isolating all cytosolic membranes.

### Global Lipid Polarity Changes in Neurons are Age-dependent and Exacerbated by AD-genes

Considering that aging and disease have a strong effect on lipid metabolism^41^, we evaluated how the lipid polarity signatures at different ages change by assessing the neutral and polar environmental fractional contributions. Supplementary Figure 4A The phasor analysis enabled us to retrieve a 3D distribution of emission spectra from all pixels in the images stained with nile red carrying information on the polarity of the environment (where x and y correspond to the polar position and z-axis, displayed as a density color map, is the frequency of pixels. Supplementary Figure 4B shows the cross-section lines drawn from the most neutral (yellow gold) to the most polar (red) possible environments. The center of mass (CM) is calculated from the frequency histogram as described in the material and methods section.

In Figure 2A we show a global whole-cell analysis where our data reveals statistically significant shifts in lipid polarity that are both age- and AD genotype-dependent. In neurons from NTg mice aging model, the overall polarity index increases in the age span, indicating a pronounced effect of age on polarity. The most substantial positive shift in polarity occurs between young and middle age (0.03 fractional units, p < 0.001; Fig. 2A, black), followed by old neurons failing to maintain the polarity and displaying a significantly mild decrease (0.01 fractional units, p < 0.001). In the 3xTg-AD model, polarity initially mirrors the NTg pattern with an early increase. However, 3xTg-AD neurons exhibit a significantly smaller increase during middle age (0.01 fractional units, p < 0.001; Fig. 2A, red) and a greater decline with aging (0.02 fractional units, p < 0.001). These results suggest that AD-related genetic alterations progressively disrupt normal age-related polarity dynamics, with effects intensifying over time. A two-way ANOVA confirmed significant effects of both age (p < 0.001) and genotype (p < 0.001) on polarity.

**Figure 2.**
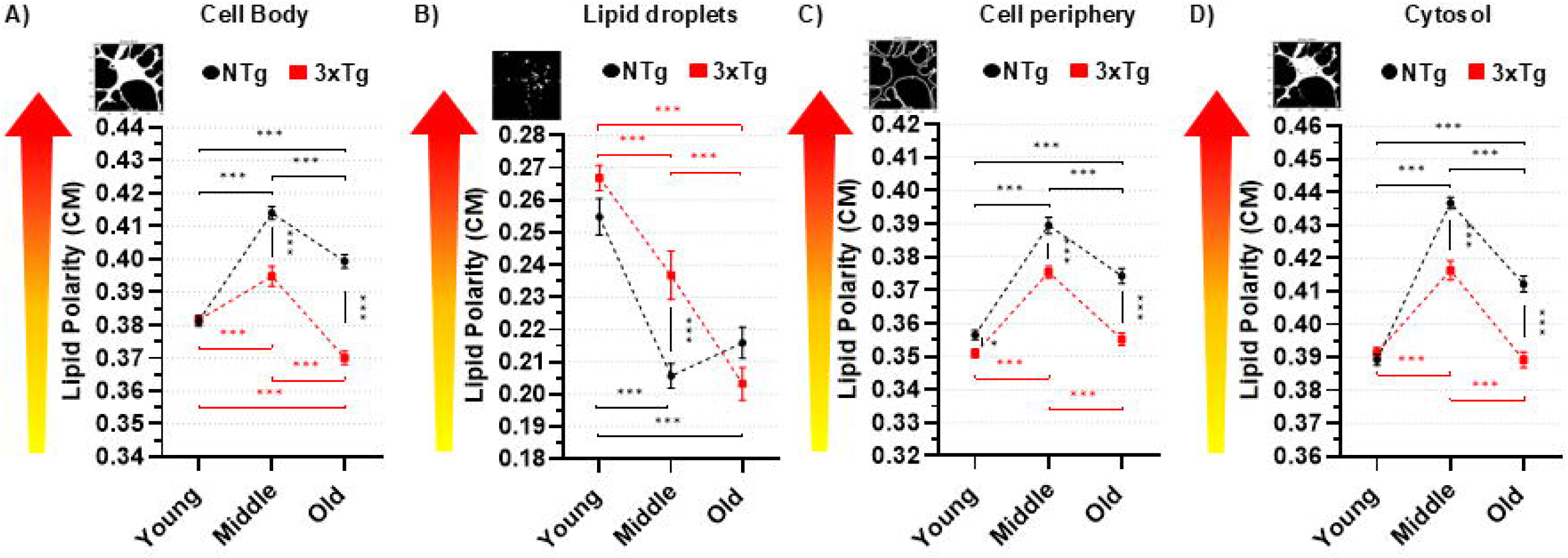
Old healthy neurons fail to maintain global lipid polarity levels, and AD-genes exacerbates such effect. Cellular compartments display similar trends with different lipid polarity signatures. A) Average Center of Mass of the global lipid polarity index for all whole cells. 8) Average Center of Mass of the global lipid polarity index for all lipid droplets in all cells. C) Average Center of Mass of the global lipid polarity index for the cell periphery. D) Average Center of Mass of the global lipid polarity index for the cytosol without lipid droplets and cell periphery. NTg corresponds to Non-transgenic neurons, while 3xTg represents triple-transgenic AD neurons. Statistical comparisons are presented as horizontal bars for age-wise comparisons, vertical bars for age-paired genotype comparisons. Error bars represent Standard Error Means (SEM). Only statistically significant p-values are shown, *** is a p-value < 0.001,** is a p-value = 0.01, * is a p-value = 0.05.

### Aging and Alzheimer’s Genotype Reduce Lipid Droplet Polarity, Increasing Neutral Lipid Content

The evaluation of the nile red emission in whole hippocampal neurons enabled us to determine the global impact of polarity as a function of aging in both NTg and 3xTg-AD genotypes. To determine whether the global decrease in polarity is mainly influenced by LDs, we conducted further analysis, as LDs are rich in neutral lipids including cholesterol and triglycerides^20^. Phasor plots and pixel frequency pertaining to the polarity histograms for the global LD analysis across different ages and conditions are shown in Supplementary Figure 5.

As expected, LDs have significantly lower polarity values than the whole-cell, confirming that LDs contain a more neutral lipid environment. In NTg neurons, lipid droplet polarity decreases with age, displaying a sharp decline between young and middle age, with a negative change of - 0.05 fractional units (p < 0.001), and then plateau in old age (Fig. 2B, black). LDs In 3xTg-AD neurons, the lipid polarity levels decrease with age, showing a decline of -0.03 fractional units by middle age (p < 0.001) and another -0.03 fractional unit drop by old age (p < 0.001) (Fig. 2B, red). Both NTg and AD-related aging reduce LD lipid polarity, making these structures richer in neutral lipids. Interestingly, only middle-aged neurons from both genotypes show a statistically significant difference (p < 0.001). The polarity levels of 3xTg-AD LDs in middle age fall between those of young and middle-aged NTg LDs, while old-age 3xTg-AD LDs resemble the polarity levels of NTg middle-aged LDs. A two-way ANOVA confirmed that age has a highly significant effect on LD polarity values (p < 0.001), while genotype also plays a contributing role (p = 0.02). However, 3xTg-AD LDs never reach lower polarity values than their NTg age-matched counterparts, suggesting that any global shifts, at the whole cell level, toward a more neutral lipid environment in neurons may not be driven by LD lipid composition, but by number fraction and size.

### Aging and AD-genotype Alter Lipid Polarity Dynamics at the Neuronal Cell Periphery

Analysis of cell periphery phasor plots initially revealed similar phasor distributions across age groups. However, examination of polarity frequency histograms elucidated subtle shifts in polarity values (Supplementary Figure 7). In NTg neurons, the periphery’s polarity substantially increases from young to middle age with a positive delta of NTg-PM-Δ_young-middle_ = 0.03 fractional units (p < 0.001), followed by mild decrease in old age with a negative delta of NTg-PM-Δ_middle-old_ = -0.01 fractional units (p < 0.001) (Fig. 2C, black). Compared to NTg, The polarity at the cell periphery in 3xTg neurons, increases less from young to middle age with a positive delta of 3xTg-PM-Δ_young-middle_ = 0.02 fractional units, (p < 0.001), followed by a decrease in polarity of the same magnitude with a negative delta of 3xTg-PM-Δ_middle-old_ = -0.02 fractional units, (p < 0.001), resulting no difference between young and old peripheries in 3xTg-AD neurons (Fig. 2C, red). Compared to other compartments, the cell periphery displays genotype differences in polarity as early as young age, this difference is minimal but significant (Δ_NTg-3xTg_ = 0.005 fractional units, p < 0.001). This difference tripled by middle age and remained constant through old age (with Δ_NTg-3xTg_ = 0.015 fractional units, p < 0.001). These findings are consistent with the expected high polarity of lipids in the plasma membrane and nearby structures, which help insulate the cell from its aqueous surroundings ^43^. They emphasize the need to consider subcellular compartments when analyzing lipid polarity and reveal that membrane lipid composition is dynamic across aging. Notably, the lipid profile at the neuronal periphery is altered by both aging and AD-associated transgenes, with significant changes detectable as early as the young age stage. This suggests a complex interaction between genetic risk for Alzheimer’s disease gene expression and aging in shaping membrane lipid composition (two-way ANOVA: Genotype p < 0.001, Age p < 0.001).

### Cytosolic Lipid Polarity Reveals Age- and AD-Related Changes Independent of Lipid Droplets

The cytosolic membranes display a higher polarity than the whole cell; this is expected as the most neutral regions of the cell, LDs and cell periphery are excluded, thus, the remaining polarity changes reflect the lipid dynamics of the rest of organelles in the cytosol (Supplementary Figure 8). Fig. 2D shows a similar pattern as the whole cell and periphery. NTg young neurons start with a relatively lower polarity that increases from young to middle age, displaying a positive delta of NTg-Cyto-Δ_young-middle_ = 0.04 fractional units (p < 0.001), this is the largest positive delta throughout the cell, followed by a polarity decrease by old age with a negative delta of NTg-Cyto-Δ_young-middle_ = -0.025 fractional units (p < 0.001) (Fig. 2D, black). In 3xTg-AD neurons, the cytosolic polarity values increase from young to middle age by a positive delta of 3xTg-Cyto-Δ_young-middle_ = 0.02 fractional units (p < 0.001), followed by a decrease of the same magnitude at old age, displaying a negative delta of 3xTg-Cyto-Δ_middle-old_ = -0.02 fractional units (p < 0.001) (Fig. 2D, red).

The above observations showing the cytosolic membranes and cell periphery displaying similar patterns of polarity changes during aging and AD-like genotype, suggest that lipid polarity alterations are reflected throughout the entire cell in proportional ways, with the exception of LDs, where the main cellular activities related to lipid metabolism and storage show more complex dynamics that we address in the following section.

### Aging and AD genotype Alter Lipid Droplet Count in Hippocampal Neurons

Lipid droplets are a storage organelle with important participation in lipid and energy homeostasis, with functions ranging from aiding in cellular metabolism and nutrient availability to stress response, many of these processes result in growth, shrinkage, fusion and fission of LDs^44^. Thus, characterizing as many features of LD as possible is an important yet challenging task to better understand their dynamics within the cell.

In this work, we have described a lipid polarity global analysis of bulk LDs in hippocampal neurons, to complement this data to characterize the LDs dynamics during aging and disease. We used the Phasor analysis of Local Image Correlation Spectroscopy (PLICS) to retrieve the size and number of LDs. From an original spectral image, we isolate the channels that contain the strongest signals for LDs. We proceed to project those channels into a 2D image which is passed to the PLICS algorithm. As a result, we obtain the average number of detected LDs and their respective size for each image. Here we present the global results from such analysis, note we only present average quantifications per cell, except when specified. Figure 3A shows the average count of LDs per neuron was similar for young neurons, with no statistical difference, between NTg and 3xTg-AD neurons averaging 22 ± 2 and 26 ± 2 LDs/optical section/cell, respectively. By middle age, both genotypes NTg and 3xTg-AD, increase their LDs count by ∼2.5 fold, now displaying an average of 55 ± 2 LDs (p_young-middle_ < 0.001) and 59 ± 2 LDs (p_young-middle_ < 0.001) per neuron, respectively, with no difference between genotypes. In old age, LD count does show a significant difference between genotypes (p_NTg-3xTg_ < 0.001) as NTg neurons show a ∼0.5-fold decrease to 29 ± 3 LDs (p_young-middle_ < 0.001), while 3xTg-AD showed a mild not statistically significant decrease to 55 ± 2 LDs. These results highlight old age as a time when LD count in neurons diverges, as both age (p < 0.001) and genotype (p < 0.001) have extremely significant effects on these results according to a 2-way ANOVA analysis.

**Figure 3.**
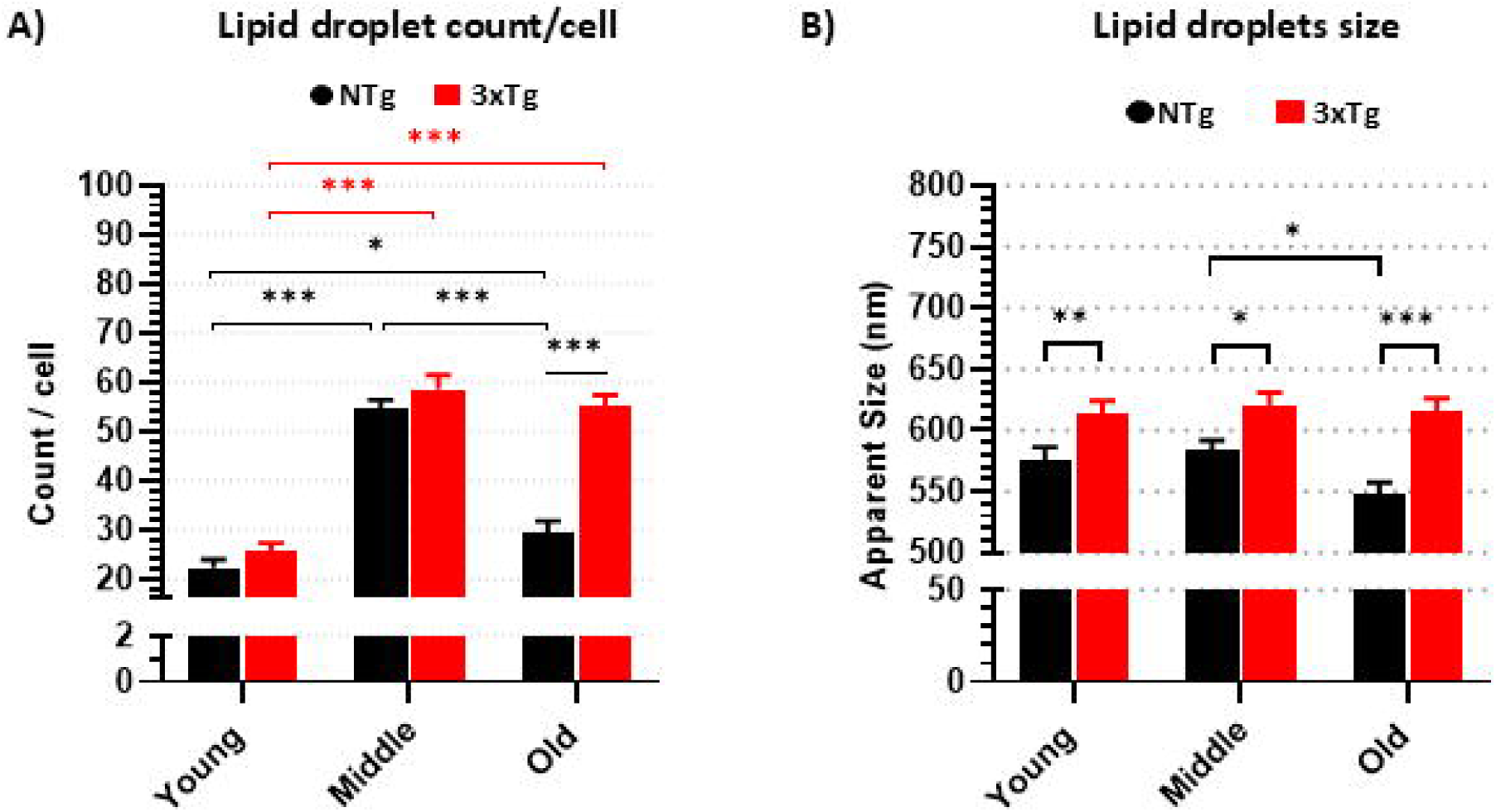
Lipid droplet count per cell and size are genotype• and age-dependent. A) Average lipid droplet count in neurons. 8) Average lipid droplet size per cell. Error bars represent Sum of Error Means (SEM). Only statistically significant p-values are shown, ***, is a p-value < 0.001, ** is a p-value = 0.01, * is a p-value = 0.05.

### Aging and AD-genotype Affect Lipid Droplet Size

Complementary to the average LD count per cell, we determined the average LD size per optical section per neuron. Figure 3B shows no statistical difference between young (576 ± 11 nm) and middle (583 ± 8 nm) ages LD size, in NTg neurons but the average size drops to 547 ± 10 nm (p = 0.03) in old age. On the other hand, 3xTg-AD neurons display no significantly different average LD sizes during aging, with sizes of 613 ± 11 nm at young age, 620 ± 11 nm at middle age, and 616 ± 10 nm at old age. However, 3xTg-AD LD sizes are consistently larger than their NTg age-pair counterparts, at young age (p _NTg-3xTg_ = 0.01), middle age (p _NTg-3xTg_ = 0.03) and old age (p _NTg-3xTg_ < 0.001). In other words, 3xTg-AD neurons fail to maintain the modest LD size present in NTg neurons. in the present work we limit ourselves to report averaged results in the form of global analysis.

### Principal Component Analysis Confirms Compartment Coordinated Age- and Gene-dependent Changes in Lipid Polarity

To better correlate global lipid polarity changes across interconnected neuronal compartments, we applied principal component analysis (PCA) to our dataset, which included all age groups and both genotypes (NTg and 3xTg-AD). PCA reduced the complexity of the data into seven principal components (PCs), of which PC1 (30%), PC2 (26%), and PC3 (15%) were selected based on Parallel Analysis, collectively explaining 72% of the total variance (Supplementary Figure 9A). Variables included genotype, age, cytosolic polarity, plasma membrane (PM) polarity, LD polarity, average count, and size. PC1 was highly correlated with cytosolic (0.83) and PM polarity (0.86), and negatively with genotype (-0.58), indicating that 3xTg neurons exhibit lower polarity across compartments (Supplementary Table 1). PC2 captured LD characteristics, with strong positive correlations with LD polarity (0.57) and size (0.69), and negative correlations with LD count (-0.66) and age (-0.67). PC3 revealed genotype-dependent polarity distribution differences, negatively correlating with LD polarity (-0.60), cytosolic polarity (-0.40), PM polarity (-0.30), and genotype (-0.58).

Age-wise and genotype-wise PCA score comparisons further support these patterns. A biplot displaying the distribution along the PC1 and PC2 scores allows us to visualize how each parameter has more or less contribution to the variance for every experimental condition (Figure 4A). In NTg neurons, PC1 scores increased significantly with age (p < 0.001), indicating a global rise in polarity across compartments (Figure 4B, black). In contrast, 3xTg neurons exhibited significantly lower PC1 scores at all ages (p < 0.001), suggesting a persistent reduction in polarity and a failure to undergo the age-related polarity increase seen in NTg neurons (Figure 4B, red).

**Figure 4.**
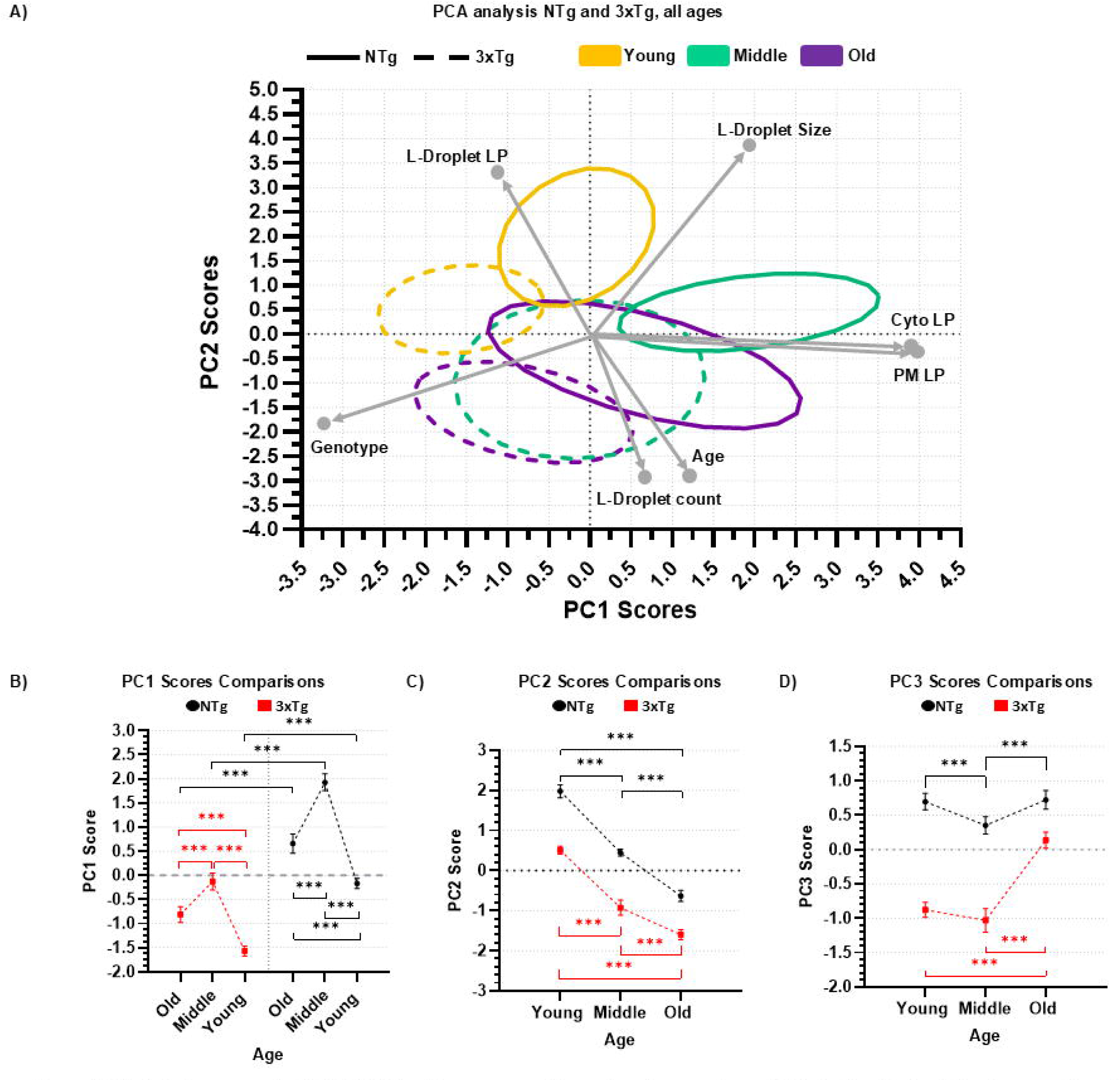
Principal Component Analysis (PCA) identifies genotype effects where the cytosolic and lipid droplet global polarity values are the main drivers of cell lipid polarity during aging. A) Confidence ellipse of PCAbiplot of all NTg and 3xTg images for a genotype comparison, each ellipse correspond to a 70% confidence interval of the data. Solid circles with arrows pointing at them correspond to the magnitude of the PC loading. B) Genotype wise comparison of PC1 scores. C) PCA biplot of NTg images only for age wise analysis. D) PCA biplot of 3xTg images only for age wise analysis. E) Age wise comparison of PC1 scores for both NTg and 3xTg images. F) Age wise comparison of PC2 scores for both NTg and 3xTg images. For every PCA analysis, magenta circles represent the loading coordinate corresponding to each cell compartment, while magenta arrows originating from the (0,0) coordinate and pointing to each circle loading representing the magnitude of the former. NTg correspond to Non• transgenic neurons, while 3xTg represents triple-transgenic AD neurons. Statistical comparisons are presented as follows, horizontal bars correspond to age-wise comparisons, vertical bar correspond to age-paired genotype comparisons. Error bars represent the mean and 95% confidence interval of the data. Only statistically significant p-values are shown, *** is a p-value < 0.001, ** is a p-value = 0.01, * is a p-value = 0.05.

PC2 scores decreased with age in NTg neurons (p < 0.001), reflecting a shift from many small LDs to fewer, larger ones with altered polarity (Figure 4C, black). 3xTg neurons displayed reduced PC2 scores early in life and continued to decline with age (p < 0.001), consistent with early and accelerated LD deterioration (Figure 4C, red).

PC3 scores in NTg neurons showed a drop in middle age but recovered by old age, while 3xTg neurons exhibited persistently negative scores in young and middle age, indicating sustained alterations in polarity distribution that only begin to normalize in old age (Figure 4D).

In summary, PC1 captures the age-related rise in polarity levels observed in NTg but disrupted in 3xTg-AD neurons, PC2 reflects structural reorganization of LDs with earlier onset in 3xTg neurons, and PC3 highlights genotype-driven differences in how polarity is distributed among compartments. These PCA-derived components reveal key structural and biochemical alterations in hippocampal neurons during aging and in the context of AD pathology.

## DISCUSSION

### Aging and AD-genes Impact Hippocampal Neurons Lipid Polarity Dynamics

Our study reveals distinct patterns of lipid polarity and lipid droplet characteristics in hippocampal neurons during aging and in the context of AD progression. The age-dependent changes in lipid polarity and lipid droplet phenotypes observed in NTg neurons contrast sharply with the disrupted lipid homeostasis seen in AD neurons. This dysregulated lipid homeostasis has been associated with aging and as a risk factor for AD ^45–47^. These findings provide new insights into the complex interplay between lipid polarity, aging and neurodegeneration processes.

The observed increase in global lipid polarity in NTg aging neurons, followed by a slight decrease in old age, suggest a dynamic regulation of lipid composition throughout the lifespan, a similar trend was observed by Díaz et al. ^48^ where the total neutral lipids concentration was decreased, and total polar lipids increased in old postmortem human brains compared to young. These age-dependent disturbances to the lipid homeostasis have been found to arise as early as middle-age and are associated with oxidative stress during normal aging with exacerbation of such effects during pathogenic neurodegeneration ^48–50^.

In contrast, AD-genotype neurons failed to show this typical age-related increase in lipid polarity, instead demonstrating an overall decrease in lipid polarity, that suggests an increase in the ratio of neutral lipids during aging. This is consistent with reports of cholesterol accumulation, which accelerates amyloid metabolism and tau phosphorylation, hallmarks of the disease ^51–53^. This disruption in lipid polarity homeostasis could contribute to the pathophysiology of AD by exacerbating alterations of membrane properties and cellular functions dependent on lipid organization ^51,54^. Our results show distinct lipid polarity signatures in different cellular compartments (cytosolic membranes, cell periphery, and lipid droplets) suggesting heterogeneity of lipid environments within neurons. Similar trends observed in cytosolic membranes and the cell periphery suggest that lipid polarity alterations are reflected throughout the entire cell with the exception of lipid droplets. This compartment-specific regulation of lipid polarity pinpoints the significance of organelle function and inter-organelle communication during aging and disease progression, as seen in organelle communication through membrane contact sites ^55,56^.

### Lipid Droplet Turnover in Hippocampal Neurons is Disrupted by Age and AD-genes

The analysis of lipid droplet count and size distribution provides valuable insights into the metabolic and stress adaptations occurring in neurons during aging and AD progression. The observed increase in lipid droplet count in middle-aged neurons of both genotypes, followed by a sustained increase in lipid droplets in the AD-genotype, but a decrease in NTg hippocampal neurons could represent a lagging indicator of neuroprotection, as observed in glia challenged to reactive oxygen species (ROS) which in order to return to normal metabolism will steadily deplete their LDs ^57,58^, highlighting an adaptive response to changing metabolic demands and increased age-related ROS production in normal aging and neuropathology. Thus, lipid droplet features during AD aging suggests dysregulation and impairment of lipid utilization and distribution, potentially contributing to increased cellular stress and dysfunction. These lipid droplet features have been observed during human aging, as microglia display an increase in size and number of lipid droplets ^59^, and in hippocampal regions, microglia accumulate lipid droplets rich in glycolipids as a response to increased ROS and inflammation ^60^. While, during normal aging lipid droplets help mitigate cholesterol toxicity through enzymatic conversion to cholesteryl esters, this process becomes disrupted during AD progression due to lysosomal lipid metabolism impairments ^61^, resulting in a further accumulation of more numerous and larger lipid droplets by microglia that also display upregulation of genes involved in lipid synthesis and biogenesis, alongside elevated ROS levels ^60,62^.

The importance of Lipid droplets during aging becomes more relevant as they help sequester toxic polyunsaturated fatty acids (PUFAs), which are prone to peroxidation during oxidative stress, these PUFAs are replaced on membranes and stored as triacylglycerol in Lipid droplets ^18,20^. This mechanism safeguards membranes from oxidative damage, particularly in hyperactive neurons that transfer excess PUFAs to astrocytic Lipid droplets via ApoE-positive particles. These lipids can be degraded through autophagy or stored as neutral lipids ^63^.

Lipid droplet biogenesis and degradation are tightly regulated to prevent lipid toxicity. Lipolysis preferentially targets large Lipid droplets, while smaller Lipid droplets accumulate under lysosomal inhibition, indicating distinct pathways for lipid turnover ^23^. Loss of oxidative phosphorylation (OXPHOS) in astrocytes further disrupts lipid homeostasis, leading to excessive free fatty acids (FFAs) and Lipid droplet accumulation ^64^. In such scenarios, neurons compensate by storing excess triacylglycerol in their own Lipid droplets ^65^.

### Lipid Organization Across Neuronal Compartments is Age- and AD-genotype dependent

The PCA results reveal interrelated changes in lipid polarity and lipid droplet characteristics across different cellular compartments. These components collectively demonstrate the complex nature of lipid alterations in aging and AD. AD-neurons consistently show reduced PC1 scores across all ages compared to normal neurons, indicating a persistent age and AD gene-driven disruption of lipid polarity homeostasis interfering with normal age-related adaptations in lipid composition and organization, such as human ApoE4 gene disturbing fatty acid transport and leading to lipid accumulation in both astrocyte and neurons in the hippocampus^65,66^. Our results show an age-related regulation of lipid droplets in both normal and AD-transgenic neurons, with AD-neurons exhibiting accelerated and more pronounced changes in lipid droplet properties as early as young ages, these early alterations in lipid storage and utilization may contribute to cellular dysfunction in AD, as they are involved in crucial lipid homeostasis throughout the cell ^18,45^. PC2, captures and age- and genotype-dependent changes in lipid droplet properties, our observations where lipid droplets increase in number and size during aging AD-neurons, aligns with other reports where microglia and astrocytes display lipid droplet enlargement, accumulation, and lipid content changes resulting from aging and disease ^45,60,71^. PC3 captures genotype-dependent differences in lipid polarity distribution, suggesting that AD-associated genetic modifications cause distinct patterns of lipid partitioning across cellular compartments, as seen in other studies where AD-associated genes impact multiple lipid pathways affecting several cellular organelles ^67^. Interestingly, these PC3 differences become less pronounced in old age, suggesting that aging is a stronger factor than the AD-genotype for altered lipid homeostasis.

### Limitations

Our study provides valuable insights into lipid polarity and lipid droplet features in aging and AD, but some limitations exist. The use of a single fluorescent probe (Nile Red) restricts our ability to distinguish between specific lipid subspecies ^45^. Future studies could employ multiple lipid-specific probes or lipidomic approaches to provide more detailed information on lipid composition changes associated with aging and AD. Our study was not sufficiently powered to reach any gender-based conclusions which are known to be an important factor in aging- and AD-development. Thus, further studies focusing on gender differences are needed ^48,68^.

## CONCLUSION

Our study has demonstrated that both normal aging and Alzheimer’s disease (AD)–related genes profoundly alter lipid polarity and lipid droplet (LD) dynamics in hippocampal neurons. In nontransgenic (NTg) neurons, aging is accompanied by a transient rise in global lipid polarity (peaking in middle age) followed by a slight decline in older animals, consistent with prior observations of increased polar lipid fractions and decreased neutral lipid stores in aged brains ^48,49^. By contrast, AD-transgenic neurons fail to exhibit this typical age dependent increase in polarity; instead, they display an overall decline in polarity—reflecting accelerated neutral lipid (e.g., cholesterol and triglycerides) accumulation—across all ages. This shift aligns with established links between cholesterol buildup, aberrant amyloid processing, and tau phosphorylation in AD pathogenesis ^51–53^. This further supports the “aging-switch” hypothesis, in which aging triggers Alzheimer’s degeneration ^69^.

Examining LD count and size distribution revealed that both NTg and AD-genotype neurons mount an early, middle-age increase in LD number, likely an adaptive response to rising oxidative stress and altered energy demands. However, whereas NTg neurons eventually cleared this increased LD load, mirroring protective LD turnover observed in glia under reactive oxygen species challenge ^57,58^, 3xTg neurons retained and even expanded their LD population into later ages. This persistent LD accumulation suggests dysregulated lipid utilization and storage in the AD-genotype context, potentially exacerbating neuronal stress. Such aberrant LD dynamics has been observed in human microglia, where aging and AD-associated lysosomal impairments drive excessive LD biogenesis, upregulation of lipid synthesis genes, and elevated ROS^60,62^.

Our principal component analysis (PCA) further highlighted that lipid polarity changes are not uniform throughout the neuron. AD-transgenic cells consistently showed lower PC1 scores, indicative of disrupted polarity homeostasis across the whole cell, in cytosolic membranes, cell periphery, and LDs, even at young ages, pointing to an early, genotype-driven breakdown of lipid regulatory networks ^65,66^. PC2 primarily captures age-related changes in LD structural features, revealing a shift toward fewer but larger LDs with altered polarity profiles in older neurons indicating progressive structural remodeling of LDs over time ^45,70^. Meanwhile, PC3 reflects genotype-dependent differences in lipid polarity distribution across compartments, consistent with previous findings that AD-related genes modulate multiple lipid pathways across diverse organelles ^67^. Interestingly, these genotype-associated differences become less pronounced in old age, suggesting that chronological aging eventually overrides genetic effects on lipid homeostasis.

Together, these findings highlight the importance of lipid polarity and droplet turnover in maintaining normal neuronal physiology and how their disruption by aging and AD genes contributes to neurodegeneration. The distinct age- and genotype-dependent patterns in lipid composition and organization across both membranes and lipid droplets, underscores the need to study lipid regulation in a compartment-specific context. The progression toward fewer, smaller, and more neutral LDs in aged neurons, and the opposite trend in AD-genotype, points to impaired lipid recycling and increased vulnerability to oxidative damage. These imbalances may compromise membrane fluidity, synaptic signaling, and metabolic resilience. Future directions should focus on the molecular pathways that regulate lipid trafficking, droplet biogenesis, and degradation, particularly at organelle contact sites, and how their dysfunction contributes to AD pathogenesis. Targeting these processes to restore balanced lipid polarity and turnover may provide new avenues for therapeutic intervention aimed at preserving neuronal function in aging and neurodegenerative disease.

## STAR METHODS

STAR ✎ METHODS
| REAGENT or RESOURCE | SOURCE | IDENTIFIER |
| --- | --- | --- |
| **Antibodies** |  |  |
| N/A |  |  |
| **Bacterial and virus strains** |  |  |
| N/A |  |  |
| **Biological samples** |  |  |
| N/A |  |  |
| **Chemicals, peptides, and recombinant proteins** |  |  |
| isoflurane, USP | Piramal Critical Care Inc | NDC Code(s): 66794-013-25 |
| Papain, Lyophilized | Worthington Biochemical | Cat #: LS003119 |
| Mouse FGF-basic (FGF-2/bFGF) Recombinant Protein | Thermo Fisher | Cat #: PMG0035 |
| Mouse FGF-basic (FGF-2/bFGF) Recombinant Protein | PeproTech | Cat #: 450-33-50UG |
| Mouse PDGF-BB Recombinant Protein | Thermo Fisher | Cat #: PMG0045 |
| Mouse PDGF-BB Recombinant Protein | PeproTech | Cat #: 315-18-10UG |
| Zeocin™ Selection Reagent | Thermo Fisher | Cat #: R25001 |
| Geneticin™ Selective Antibiotic (G418 Sulfate) (50 mg/mL) | Thermo Fisher | Cat #: 10131035 |
| Nile Red, 9-(diethylamino)-5H-benzo[a]phenoxazin-5-one | Cayman Chemical | Cat #: 30787 |
| Dimethyl sulfoxide DMSO | Sigma-Aldrich | Cat #: D2650-100ML |
| LipiBlue | Dojindo | Cat #: LD01 |
| LysoTracker™ Deep Red | Thermo Fisher | Cat #: L12492 |
| FM™ 1-43 Dye (N-(3-Triethylammoniumpropyl)-4-(4-(Dibutylamino) Styryl) Pyridinium Dibromide) | Thermo Fisher | Cat #: T3163 |
| **Critical commercial assays** |  |  |
| N/A |  |  |
| **Deposited data** |  |  |
| Raw and analyzed data | This paper | Upon request |
| Original spectral images | This paper | Upon request |
| **Experimental models: Cell lines** |  |  |
| Human: SH-SY5Y | ATCC | Cat #: CRL-2266 |
| Mouse: Hippocampal Neurons | This paper | N/A |
| **Experimental models: Organisms/strains** |  |  |
| Mouse: NTg: C57/BL6 | Charles River | Strain 027 |
| Mouse: 3xTg B6;129-Psen1tm1Mpm Tg(APP<sup>Swe</sup>,tauP301L)1Lfa/Mmjax | The Jackson Lab | JAX MMRRC Stock# 034830 |
| **Oligonucleotides** |  |  |
| N/A |  |  |
| **Recombinant DNA** |  |  |
| N/A |  |  |
| **Software and algorithms** |  |  |
| Python 3.11.9 | Python Software Foundation | www.python.org |
| GraphPad Prism Version 10.4.1 | GraphPad Software | https://www.graphpad.com |
| 2-Component Spectral Phasors | This paper | https://github.com/ABBMresearch/2Component_SpectralIPhasors |
| PLICS algorithm Python version | This paper | https://github.com/ABBMresearch/PLICS_python_ver1 |
| PLICS algorithm Original MATLAB version | Scipioni et al. 2016 |  |
| **Other** |  |  |
| Rodent food: Teklad global soy protein-free extruded | Envigo Teklad | Cat #: 2020X |
| Cover Glasses, Squares, 15 mm, Thickness 0.09-0.12 mm | Carolina Biologicals | Cat #: 633071 |
| BrainBits HIBERNATE A WITHOUT CALCIUM | BrainBits HACA500 | Cat #: NC0176976 |
| BrainPhys™ Neuronal Medium and SM1 Kit | StemCell Technologies | Cat #: 05790 |
| Neurobasal A | Thermo Fisher | Cat #: 10888022 |
| B27 supplement | Thermo Fisher | Cat #: 17504044 |
| GlutaMAX™ Supplement | Thermo Fisher | Cat #: 35050-061 |
| Optiprep | Cosmo Bio | Cat #: AXS-1114542 |
| Poly-D-lysine | Sigma-Aldrich | Cat #: P6407-5MG |
| B-27™ Plus Neuronal Culture System | Thermo Fisher | Cat #: A3653401 |
| Incubator: Forma™ Series II Water-Jacketed CO2 Incubator, 184L | Thermo-Forma | Cat #: 3130 |
| GenClone 25-209, TC Treated Flasks, 250ml, Vent Growth Area: 75.0cm<sup>2</sup>, 5 per Sleeve, 100 Flasks/Unit | Genesee Scientific | Cat #: 25-209 |
| GenClone 25-503 DMEM/Ham's F-12, with L-Glutamine, with 15 mM HEPES, 6 x 500 mL/Unit | Genesee Scientific | Cat #: 25-503 |
| GenClone 25-514H GenClone™ Fetal Bovine Serum, 100% U.S. Origin, Heat Inactivated, 500 mL/Unit | Genesee Scientific | Cat #: 25-514H |
| GenClone 25-508 DPBS, 1X, without Ca, Mg, Phenol Red, 0.1um Sterile Filtered, 6 x 500 mL/Unit | Genesee Scientific | Cat #: 25-508 |
| Trypsin-EDTA (0.25%), phenol red | Thermo Fisher | Cat #: 25200056 |
| 8 Well chambered cover Glass with #1.5 high performance cover glass - 57mm x 25mm base | Cellvis | Cat #: C8-1.5H-N |

### Animal models

The animal models used in this study includes a non-transgenic control (NTg) mouse C57/BL6, bred in house but obtained originally from Charles River (San Diego, CA). We also used the Triple transgenic mouse model of AD (3xTg-AD) with human transgenes APP (Swe, KM670/671NL) and Tau (P301L) under the control of Thy1.2 promotors in a PS1 (M146V). The mice were housed 1 to 5 per cage and fed rodent diet 2020X with 24% calories from protein, 16% from fat, and 60% from carbohydrates. Room temperature was maintained at 21□C, humidity at 16–25% with a 12-h light-dark cycle. This study includes three to four mice of different ages for biological relevance. The ages (in months, mO) and genders (female, F; male, M) are as follows: Young NTg 2-5 months old: 2(F), and 2(M); 3xTg: 3(F), and 1(M); Middle-aged NTg: 9-16 months old 2(F), and 2(M); 3xTg: 2(F), and 2(M); Old NTg: 3(M); and 3xTg: 3(F).

### Primary mouse neuron culture

Isolation and culture of adult hippocampal neurons followed the procedure of Brewer et al.^72^ . Briefly, mice from Brewer’s aging colony were anesthetized with vapors from 1 mL of isoflurane in a paper towel in a beaker. Hippocampal and overlaying entorhinal cortical neurons were isolated. Hippocampus portions of each hemisphere are sliced at 0.5 mm and combined into Hibernate AB medium and placed for 8 min into a 30□C water bath. Digestion of the tissue was done with 2 mg/mL of papain in Hibernate A minus calcium and 0.5 mM Glutamax in a dry bath at 30□C with constant shaking at a speed of 125 rpm for 30 min. The tissue was then triturated and transferred to a separate 15 mL tube of Optiprep with 4 layered densities. The two gradients were centrifuged at 800 g for 15 min. The neuron enriched fractions were collected and transferred to 5 mL of Hibernate AB. The cell suspension was centrifuged twice for 1 min at 200 × *g* and the supernatant were discarded. The cells were plated onto 15 mm coverslips coated overnight with 0.1 mg/mL Poly-D-lysine in water, at a density of 50,000 cell/cm^2^ in Neurobasal A with B27 supplemented with 5 ng/mL Mouse FGF-basic Recombinant Protein and 5 ng/mL PDGF-BB for trophic support. The medium is adjusted from 270 to 290 mOsm with 5 M NaCl. One-half medium changes were made on day 3 with 10 ng/mL growth factors, assuming consumption of the prior growth factors. Another one-half medium change was made on day 6 and 10 using BrainPhys with SM1 and 10 ng/mL each of FGF-2 and PDGF-BB. The cells were cultured for 10–15 days until the cells exhibited projections and formed networks at 37□C in 5% CO_2_ and 9% O_2_ at saturated humidity prior to use for experimentation. Neurons were imaged in culture medium and 5% CO_2_ at 37 °C.

### Culture of human stable cell line

The stable human neuroblastoma cell line SH-SY5Y was maintained in culture in Nunc EasYFlask T-75 flasks using cell medium DMEM/Ham’s F-12 supplemented with L-Glutamine, 15 mM HEPES, 10% Fetal Bovine Serum, heat Inactivated and, an antibiotic cocktail of Zeocin 100 µg/mL, and Geneticin 120 µg/mL. Cells were passed every third day or when confluency reached 80% by washing the culture with Dulbecco’s PBS without Ca, Mg, Phenol Red and de-attached, with Trypsin-EDTA (0.25%) phenol red. Cultures were split 1:4 every 2 days. Cultures were always maintained at 37□C in 5% CO_2_ at saturated humidity. For experiments, a cell density of ∼30,000 cells per cm^2^ were plated onto 8-well chambered cover glass dishes.

### Fluorescent labeling and probes

Nile Red stock solution was prepared at 1 mg/mL in DMSO and further diluted to a 500 μM working solution in DMSO. Cells were incubated with nile red for one hour in 5% CO_2_ and 9% O_2_ at a final concentration of 500 nM before imaging. Lipid droplets were stained with LipiBlue using a 0.1 mM DMSO stock solution, as per the manufacturer’s instructions, and diluted 1:1000 in the cell medium for incubation. Lysosomes were labeled with LysoTracker Deep Red from a 1 mM stock solution, following the manufacturer’s guidelines, and used at a final dilution of 1:1000. Early endosomes and recycling vesicles were tracked using FM1-43, prepared as a 5 mM stock solution per the manufacturer’s instructions and applied at a 1:1000 dilution. All incubations were conducted at 37°C in 5% CO_2_ and 9% O_2_ under saturated humidity. SH-SY5Y cells were stained with two dyes at a time to assess organelle colocalization, using the following pairs: LipiBlue–Nile Red, LipiBlue–LysoTracker, and LipiBlue–FM1-43. Primary neuron cultures were incubated only with Nile Red.

### Confocal hyperspectral imaging

Hyperspectral confocal images were acquired using a Zeiss LSM 880 Axio Observer microscope (Carl Zeiss, Germany) equipped with a 63x, 1.4 NA Plan-Apochromat oil immersion objective (DIC M27). Single-plane hyperspectral images were captured in pseudo-photon counting mode using a 1 × 32 channel GaAsP spectral PMT detector, covering a spectral range of 410–695 nm with a 9 nm bandwidth per channel. Images were acquired at a resolution of 512 × 512 pixels, with a pixel size of ∼0.066 μm and a pixel dwell time of 4.10 μs. The averaging method used was a 16-line summation, with a digital gain of 1, digital offset of 0, and a pinhole size of 49 μm. Imaging was conducted at 37°C in 5% CO₂ under saturated humidity conditions. Excitation was performed using a 488 nm single-wavelength Argon-Ion laser (LASOS, Jena, Germany). For each experimental condition, 20–40 randomly selected regions of interest (ROIs) per replicate were imaged, ensuring that a single neuron was present in the field of view.

For LipiBlue colocalization analysis with other probes in SH-SY5Y cells using, confocal images were acquired using the same instrument described above, with the following acquisition parameters: image size of 512 × 512 pixels, 16-bit depth, pixel size of ∼132 µm, pixel dwell time of 8.19 µs, unidirectional scanning, and an averaging method of one frame. The pinhole size was set to 90 µm, digital gain to 1, digital offset to 0, and master gain for all PMT detectors to 700. A total of 30 randomly selected field of views per condition were imaged, with multiple cells in the field of view. Fluorescence excitation and collection parameters were as follows: LipiBlue was excited using the internal pulsed high-power 405 nm laser with a -405 beam splitter, and emission was collected between 410–480 nm. Nile Red and FM1-43 were excited with a 488 nm single-wavelength Argon-Ion laser (LASOS, Jena, Germany) using a 488 nm beam splitter. Nile red fluorescence was collected within a 525–605 nm bandwidth, while FM1-43 emission was detected between 510–700 nm. LysoTracker was excited using the internal 561 nm diode laser with a 488/561 beam splitter, and its emission was collected in the 570–650 nm range.

### Image segmentation and processing pipeline

Image segmentation and the phasor analysis were done using a customized python pipeline described as follows. Source code is available in the first author’s GitHub repository (https://github.com/ABBMresearch/2Component_SpectralPhasors & https://github.com/ABBMresearch/PLICS_python_ver1).

Compartment-specific binary masks were generated as follows. From the original 32-channel spectral image, a 2D projection of channels 29–32 was created. A user-defined threshold, initialized using the Triangle method, was then applied, converting values above the threshold into a binary image representing the whole cell body. From this mask, a 6-pixel-thick contour was drawn, extending 3 pixels inward and outward from the cell edge, defining the plasma membrane (PM) region.

For the lipid droplet binary mask, a 2D projection of spectral channels 21–22, which capture the peak emission of lipid droplets (∼ 587 to 605 nm) was generated (Lipid droplet 2D image). The autocorrelation function (ACF) of this image was calculated, and the full width at half maximum (FWHM) of the ACF center was used to determine the size of a median filter. This filter was applied to the Lipid droplet 2D image, and the filtered result was subtracted from the original, enhancing Lipid droplet features. The final Lipid droplet binary mask was obtained by applying the same user-defined thresholding method described earlier.

To create the cytosolic binary mask, logical operations were applied: first, the plasma membrane mask was subtracted from the whole-cell binary image, and then the Lipid droplet mask was subtracted from the result. This process effectively excluded the plasma membrane and Lipid droplet regions, isolating the cytosolic compartment.

For spectral phasor compartment analysis, each binary mask corresponding to a specific cellular compartment was convolved with each spectral channel of the original image, followed by a spectral phasor transformation, as detailed in the next section. The resulting phasor G and S coordinates for each pixel were then plotted in a spectral phasor plot, which represents a 3D distribution of phasors along a linear combination of neutral (gold-yellow) and polar (red) components (see Results section).

Each phasor was projected onto this linear combination to generate a unidimensional distribution, where the frequency of phasors represents the fractional intensity contribution from neutral and polar components, termed the Lipid Polarity index. Since lipid polarity frequency distributions are not always Gaussian, the center of mass was calculated to quantify shifts toward more or less polar lipid environments. The center of mass provides a weighted average of lipid polarity values for each image. This process was repeated for every image in the dataset across all experimental conditions and cellular compartments. This comprehensive approach enables the assessment of lipid polarity changes in neurons under different conditions and within distinct cellular regions.

### Spectral phasor transformation and fractional analysis

The spectral phasor transform maps the complex spectral information of an image onto a 3D phasor plot by applying Fourier cosine (Eq. 1) and sine (Eq. 2) transforms to each pixel in the hyperspectral image. This transformation results in polar coordinates (G, S). A detailed explanation of the spectral phasor method, its properties, and applications can be found in previous studies ^33–35^.

In summary, the intensity at each pixel (i, j) across the detected optical spectrum (λ*_max_*-λ*_mix_*), measured at each λ interval or bandwidth channel, is used in the sine and cosine transforms. These calculations are performed using the first harmonic (n = 1), and the resulting g and s coordinates preserve the properties of vector addition.

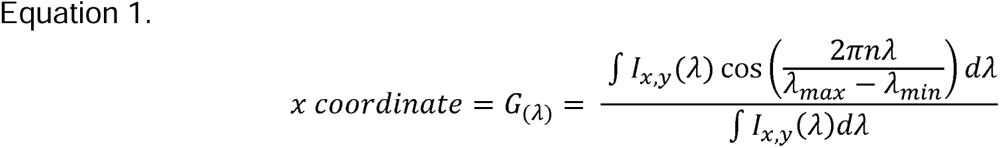

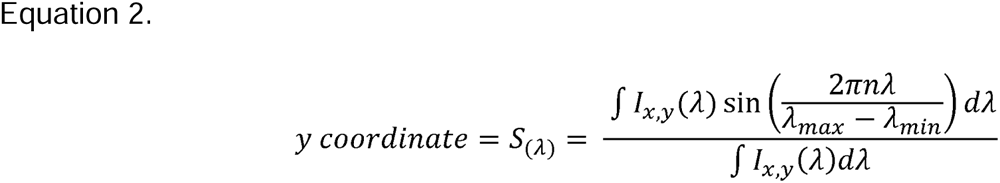

Our spectral phasor data consists of two primary components, corresponding to the two extreme spectral emissions of Nile Red in the presence of neutral (∼585 nm, yellow) and polar (∼640 nm, red) lipids, referred to as Component 1 (C1) and Component 2 (C2), respectively. These components serve as reference points to determine each phasor’s fractional position along the C1-C2 axis. Each (G, S) coordinate pair is projected onto the C1-C2 vector line using Equation 3:

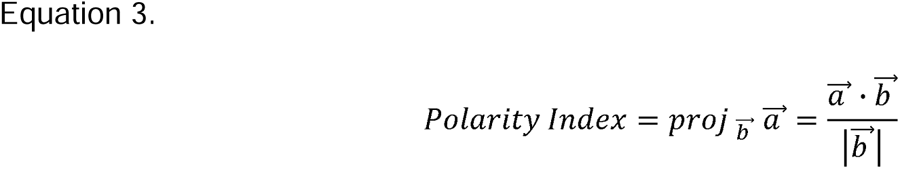

The Polarity Index is calculated by projecting vector *<u>a</u>*, (phasor (i,j)), onto vector *<u>b</u>*, (C1-C2 line), both originating from C1. This projection yields the Lipid Polarity Index for each phasor along the C1-C2 axis. The resulting Lipid Polarity Indices are then plotted as a frequency histogram, illustrating the distribution of lipid polarity across the image. To compare these distributions across conditions, the center of mass (CM) of each histogram is computed using Equation 4, providing a weighted average lipid polarity value, where *Fi* corresponds to the frequency of instances for polarity index *i*:

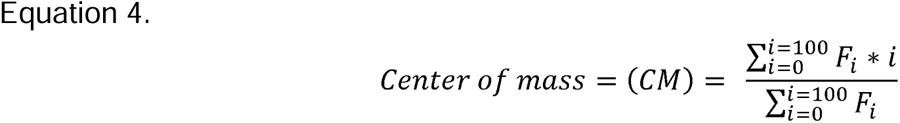

### Phasor analysis of Local Image Correlation Spectroscopy (PLICS)

Phasor Analysis of Local Image Correlation Spectroscopy (PLICS) is a fast and fit-free algorithm capable of mapping the heterogeneity of spatial correlation functions based on the phasor analysis of local ICS, in other words, the algorithm can discriminate between structures of different sizes in a heterogeneous sample with high accuracy ^24^.

Shortly, the Lipid droplet image is generated by extracting and 2D-projecting the spectral channels 21–22 from the original spectral image. We calculate the global autocorrelation function (ACF) of the Lipid droplet image, *I_x,y_*, defined as:

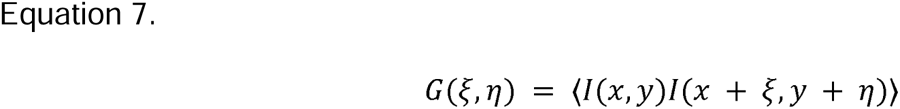

Where *G(ξ,n*) is the 2-D spatial ACF at the *ξ* and η limits, x and y respectively. *I_x,y_* is the intensity at each pixel and the brackets indicates the average is carried out on the entire image.

The spatial ACF is a normalized function containing information about the average typical width and magnitude of each spatial fluctuation in the image. These two parameters can be used to calculate the size of the fluorescent elements and their concentration, respectively, resulting in the following equation:

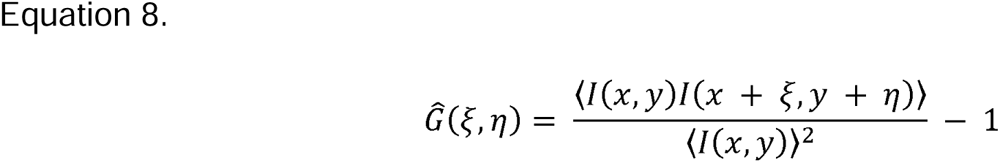

Conventional image correlation spectroscopy considers the whole image to calculate the spatial ACF, in the case of PLICS, a smaller mask of size *m x m* centered on the pixel *(i,j)* is shifted throughout the entire image using single steps along the *x* and *y* axis of the image, as a result, a 2D ACF is calculated for every *m x m* sub-image into a Local ACF matrix. Each 2D ACF is computed with a 2D Fast Fourier Transform (FFT) algorithm to accelerate the computing time.

Finally, a normalized 2D ACF is obtained for each sub image and stored in a Local ACF matrix.

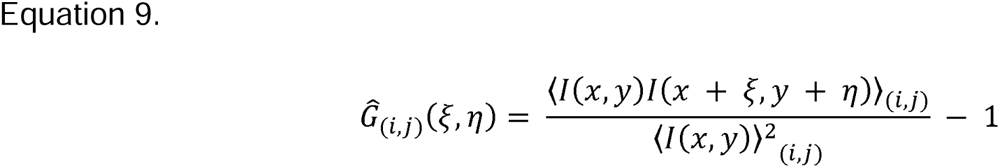

This Local ACF matrix is now converted into a 3D matrix, where the third dimension contains the ACF function *G_i,j_*(χ) obtained from every sub image of size *m x m* centered at the pixel *(i,j)*.

For each pixel of the image, that is, along the third dimension of the Local ACF matrix, two phasor coordinates *g* and *s* are calculated by means of an FFT calculation, as follows:

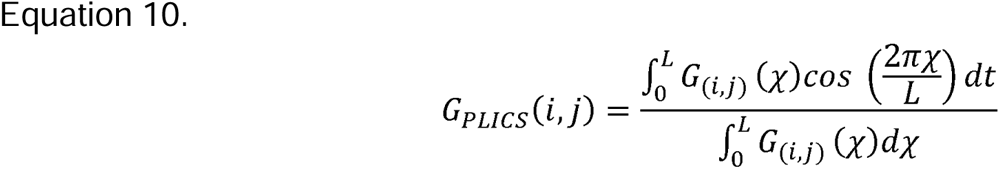

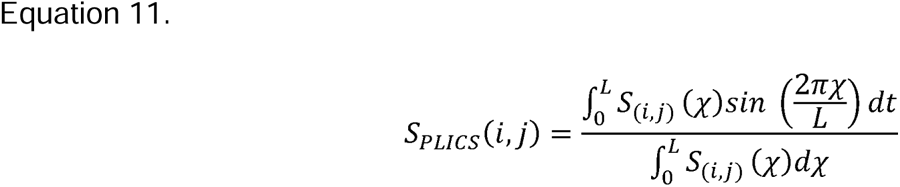

Being *L* the period of the FFT corresponding to m/2. The *G* and *S* coordinates can be plotted in a phasor plot. These phasor position contains unbiased information about the local correlation function *G_i,j_*(χ) . Followed by retrieval of the number and size of the different Local ACFs.

Given that the resolution of our imaging set up is diffraction limited, the absolute size of the segmented object is the convolution of the particle’s size and the PSF of the microscope. The minimum achievable resolution in our system is theoretically calculated using Abbe’s diffraction limit equation, ∼ 215 nm for our lateral resolution. We corroborated our analysis to effectively retrieve apparent sizes above the diffraction limit by imaging yellow fluorescent beads of 200 and 500 nm, successfully calculating apparent sizes of 200 and 500 nm.

### Statistical Analysis

All statistical analyses were performed using GraphPad Prism. Each age and genotype condition included N = 3–4 biological replicates, with 20–40 images acquired per condition, yielding a total n > 100 per condition.

Comparisons of Lipid Polarity Center of Mass (CM) values across different ages and genotypes for each cellular compartment were conducted using a two-way ANOVA with uncorrected Fisher’s LSD and a single pooled variance. The same approach was applied for comparisons of Lipid droplet average size and count, with significant p-values reported in the graphs. For Lipid droplet colocalization analysis, a Kruskal-Wallis test with Dunn’s multiple comparisons correction was used to assess differences between experimental conditions. Lipid droplet size frequency histograms were generated using a bin size of 66, corresponding to the pixel size in the original spectral images. For Principal Component Analysis (PCA), genotype values were numerically coded as 1 (NTg) and 2 (3xTg-AD), while age groups were assigned 1 (young), 2 (middle), and 3 (old). Other variables, including cytosol polarity, cell periphery polarity, Lipid droplet polarity, average size, and average count, were used without modification. All data were standardized to a mean of 0 and a standard deviation of 1. Principal components were selected based on parallel analysis with a 95% confidence interval. Differences in PC scores across ages and genotypes were assessed using a two-way ANOVA with uncorrected Fisher’s LSD and a single pooled variance. Finally, to generate confidence ellipse data, we used the Real Statistics Resource Pack software (Release 8.9.1). Copyright (2013 – 2023) Charles Zaiontz. www.real-statistics.com.

## Notes

https://github.com/ABBMresearch/2Component_SpectralPhasors

https://github.com/ABBMresearch/PLICS_python_ver1

### Competing Interest Statement

The authors have declared no competing interest.

### Funding

- Consejo Nacional de Humanidades, Ciencias y Tecnologías: 739656
- National Institutes of Health: 2P41GM103540
- National Institutes of Health: RF1 AG058218

## REFERENCES

1. Hornemann, Th. (2021). Mini review: Lipids in Peripheral Nerve Disorders. Neurosci Lett 740, 135455. 10.1016/j.neulet.2020.135455.

2. Mesa-Herrera, Taoro-González, Valdés-Baizabal, Diaz, and Marín (2019). Lipid and Lipid Raft Alteration in Aging and Neurodegenerative Diseases: A Window for the Development of New Biomarkers. Int J Mol Sci 20, 3810. 10.3390/ijms20153810.

3. Naudí, A., Cabré, R., Jové, M., Ayala, V., Gonzalo, H., Portero-Otín, M., Ferrer, I., and Pamplona, R. (2015). Lipidomics of Human Brain Aging and Alzheimer’s Disease Pathology. In International Review of Neurobiology (Academic Press Inc.), pp. 133–189. 10.1016/bs.irn.2015.05.008.

4. Stordeur, C., Puth, K., Sáenz, J.P., and Ernst, R. (2014). Crosstalk of lipid and protein homeostasis to maintain membrane function. bchm 395, 313–326. 10.1515/hsz-2013-0235.

5. Tracey, T.J., Steyn, F.J., Wolvetang, E.J., and Ngo, S.T. (2018). Neuronal Lipid Metabolism: Multiple Pathways Driving Functional Outcomes in Health and Disease. Front Mol Neurosci 11. 10.3389/fnmol.2018.00010.

6. Mota-Martorell, N., Andrés-Benito, P., Martín-Gari, M., Galo-Licona, J.D., Sol, J., Fernández-Bernal, A., Portero-Otín, M., Ferrer, I., Jove, M., and Pamplona, R. (2022). Selective brain regional changes in lipid profile with human aging. Geroscience 44, 763– 783. 10.1007/s11357-022-00527-1.

7. Söderberg, M., Edlund, C., Kristensson, K., and Dallner, G. (1990). Lipid Compositions of Different Regions of the Human Brain During Aging. J Neurochem 54, 415–423. 10.1111/j.1471-4159.1990.tb01889.x.

8. Vance, J.E. (2012). Dysregulation of cholesterol balance in the brain: Contribution to neurodegenerative diseases. Preprint, 10.1242/dmm.010124 https://doi.org/10.1242/dmm.010124.

9. Yin, F. (2023). Lipid metabolism and Alzheimer’s disease: clinical evidence, mechanistic link and therapeutic promise. FEBS J 290, 1420–1453. 10.1111/febs.16344.

10. Fitzner, D., Bader, J.M., Penkert, H., Bergner, C.G., Su, M., Weil, M.-T., Surma, M.A., Mann, M., Klose, C., and Simons, M. (2020). Cell-Type- and Brain-Region-Resolved Mouse Brain Lipidome. Cell Rep 32, 108132. 10.1016/j.celrep.2020.108132.

11. Anand, K., and Dhikav, V. (2012). Hippocampus in health and disease: An overview. Preprint, 10.4103/0972-2327.104323 https://doi.org/10.4103/0972-2327.104323.

12. Small, S.A. (2001). Age-Related Memory Decline. Arch Neurol 58. 10.1001/archneur.58.3.360.

13. Sassano, M.L., Felipe-Abrio, B., and Agostinis, P. (2022). ER-mitochondria contact sites; a multifaceted factory for Ca2+ signaling and lipid transport. Front Cell Dev Biol 10. 10.3389/fcell.2022.988014.

14. Giordano, F. (2018). Non-vesicular lipid trafficking at the endoplasmic reticulum– mitochondria interface. Biochem Soc Trans 46, 437–452. 10.1042/BST20160185.

15. Reinisch, K.M., De Camilli, P., and Melia, T.J. (2025). Lipid Dynamics at Membrane Contact Sites. Annu Rev Biochem. 10.1146/annurev-biochem-083024-122821.

16. Athenstaedt, K., and Daum, G. (2006). The life cycle of neutral lipids: Synthesis, storage and degradation. Preprint, 10.1007/s00018-006-6016-8 https://doi.org/10.1007/s00018-006-6016-8.

17. Liang, Y., and Wen, Z. (2014). Bio-based nutraceuticals from biorefining. In Advances in Biorefineries (Elsevier), pp. 596–623. 10.1533/9780857097385.2.596.

18. Ralhan, I., Chang, C.-L., Lippincott-Schwartz, J., and Ioannou, M.S. (2021). Lipid droplets in the nervous system. Journal of Cell Biology 220. 10.1083/jcb.202102136.

19. Zadoorian, A., Du, X., and Yang, H. (2023). Lipid droplet biogenesis and functions in health and disease. Nat Rev Endocrinol 19, 443–459. 10.1038/s41574-023-00845-0.

20. Danielli, M., Perne, L., Jarc Jovičić, E., and Petan, T. (2023). Lipid droplets and polyunsaturated fatty acid trafficking: Balancing life and death. Front Cell Dev Biol 11. 10.3389/fcell.2023.1104725.

21. Paar, M., Jüngst, C., Steiner, N.A., Magnes, C., Sinner, F., Kolb, D., Lass, A., Zimmermann, R., Zumbusch, A., Kohlwein, S.D., et al. (2012). Remodeling of Lipid Droplets during Lipolysis and Growth in Adipocytes. Journal of Biological Chemistry 287, 11164–11173. 10.1074/jbc.M111.316794.

22. Pu, J., Ha, C.W., Zhang, S., Jung, J.P., Huh, W.-K., and Liu, P. (2011). Interactomic study on interaction between lipid droplets and mitochondria. Protein Cell 2, 487–496. 10.1007/s13238-011-1061-y.

23. Schott, M.B., Weller, S.G., Schulze, R.J., Krueger, E.W., Drizyte-Miller, K., Casey, C.A., and McNiven, M.A. (2019). Lipid droplet size directs lipolysis and lipophagy catabolism in hepatocytes. Journal of Cell Biology 218, 3320–3335. 10.1083/jcb.201803153.

24. Scipioni, L., Gratton, E., Diaspro, A., and Lanzanò, L. (2016). Phasor Analysis of Local ICS Detects Heterogeneity in Size and Number of Intracellular Vesicles. Biophys J 111, 619–629. 10.1016/j.bpj.2016.06.029.

25. Diaz, G., Melis, M., Batetta, B., Angius, F., and Falchi, A.M. (2008). Hydrophobic characterization of intracellular lipids in situ by Nile Red red/yellow emission ratio. Micron 39, 819–824. 10.1016/j.micron.2008.01.001.

26. Han, X., Rozen, S., Boyle, S.H., Hellegers, C., Cheng, H., Burke, J.R., Welsh-Bohmer, K.A., Doraiswamy, P.M., and Kaddurah-Daouk, R. (2011). Metabolomics in Early Alzheimer’s Disease: Identification of Altered Plasma Sphingolipidome Using Shotgun Lipidomics. PLoS One 6, e21643. 10.1371/journal.pone.0021643.

27. Raz, N., Lindenberger, U., Rodrigue, K.M., Kennedy, K.M., Head, D., Williamson, A., Dahle, C., Gerstorf, D., and Acker, J.D. (2005). Regional Brain Changes in Aging Healthy Adults: General Trends, Individual Differences and Modifiers. Cerebral Cortex 15, 1676– 1689. 10.1093/cercor/bhi044.

28. Varma, V.R., Büşra Lüleci, H., Oommen, A.M., Varma, S., Blackshear, C.T., Griswold, M.E., An, Y., Roberts, J.A., O’Brien, R., Pletnikova, O., et al. (2021). Abnormal brain cholesterol homeostasis in Alzheimer’s disease—a targeted metabolomic and transcriptomic study. NPJ Aging Mech Dis 7. 10.1038/s41514-021-00064-9.

29. Xiong, H., Callaghan, D., Jones, A., Walker, D.G., Lue, L.-F., Beach, T.G., Sue, L.I., Woulfe, J., Xu, H., Stanimirovic, D.B., et al. (2008). Cholesterol retention in Alzheimer’s brain is responsible for high β- and γ-secretase activities and Aβ production. Neurobiol Dis 29, 422–437. 10.1016/j.nbd.2007.10.005.

30. Greenspan, P., Mayer, E.P., and Fowler, S.D. (1985). Nile red: a selective fluorescent stain for intracellular lipid droplets. J Cell Biol 100, 965–973. 10.1083/jcb.100.3.965.

31. Sackett, D.L., and Wolff, J. (1987). Nile red as a polarity-sensitive fluorescent probe of hydrophobic protein surfaces. Anal Biochem 167, 228–234. 10.1016/0003-2697(87)90157-6.

32. Digman, M.A., Caiolfa, V.R., Zamai, M., and Gratton, E. (2008). The phasor approach to fluorescence lifetime imaging analysis. Biophys J 94. 10.1529/biophysj.107.120154.

33. Malacrida, L. (2023). Phasor plots and the future of spectral and lifetime imaging. Preprint at Nature Research, 10.1038/s41592-023-01906-y https://doi.org/10.1038/s41592-023-01906-y.

34. Malacrida, L., Ranjit, S., Jameson, D.M., and Gratton, E. (2021). The Phasor Plot: A Universal Circle to Advance Fluorescence Lifetime Analysis and Interpretation. 10.1146/annurev-biophys-062920.

35. Fereidouni, F., Bader, A.N., and Gerritsen, H.C. (2012). Spectral phasor analysis allows rapid and reliable unmixing of fluorescence microscopy spectral images. Opt Express 20, 12729. 10.1364/OE.20.012729.

36. Gajo, C., Shchepanovska, D., Jones, J.F., Karras, G., Malakar, P., Greetham, G.M., Hawkins, O.A., Jordan, C.J.C., Curchod, B.F.E., and Oliver, T.A.A. (2024). Nile Red Fluorescence: Where’s the Twist? J Phys Chem B 128, 11768–11775. 10.1021/acs.jpcb.4c06048.

37. Roy, D., and Tedeschi, A. (2021). The Role of Lipids, Lipid Metabolism and Ectopic Lipid Accumulation in Axon Growth, Regeneration and Repair after CNS Injury and Disease. Cells 10, 1078. 10.3390/cells10051078.

38. Krishna, M.M.G. (1999). Excited-State Kinetics of the Hydrophobic Probe Nile Red in Membranes and Micelles. J Phys Chem A 103, 3589–3595. 10.1021/jp984620m.

39. Tatenaka, Y., Kato, H., Ishiyama, M., Sasamoto, K., Shiga, M., Nishitoh, H., and Ueno, Y. (2019). Monitoring Lipid Droplet Dynamics in Living Cells by Using Fluorescent Probes. Biochemistry 58, 499–503. 10.1021/acs.biochem.8b01071.

40. Minò, A., Cinelli, G., Lopez, F., and Ambrosone, L. (2023). Optical Behavior of Nile Red in Organic and Aqueous Media Environments. Applied Sciences 13, 638. 10.3390/app13010638.

41. Cuenca-Bermejo, L., Prinetti, A., Kublickiene, K., Raparelli, V., Kautzky-Willer, A., Norris, C.M., Pilote, L., and Herrero, M.T. (2023). Fundamental neurochemistry review: Old brain stories - Influence of age and sex on the neurodegeneration-associated lipid changes. J Neurochem 166, 427–452. 10.1111/jnc.15834.

42. Malacrida, L., and Gratton, E. (2018). LAURDAN fluorescence and phasor plots reveal the effects of a H2O2 bolus in NIH-3T3 fibroblast membranes dynamics and hydration. Free Radic Biol Med 128, 144–156. 10.1016/j.freeradbiomed.2018.06.004.

43. Bruce Alberts, Alexander Johnson, Julian Lewis, Martin Raff, Keith Roberts, and Peter Walter. (2002). Molecular Biology of the Cell 4th edition. (New York: Garland Science).

44. Olzmann, J.A., and Carvalho, P. (2019). Dynamics and functions of lipid droplets. Nat Rev Mol Cell Biol 20, 137–155. 10.1038/s41580-018-0085-z.

45. Cerasuolo, M., Di Meo, I., Auriemma, M.C., Paolisso, G., Papa, M., and Rizzo, M.R. (2024). Exploring the Dynamic Changes of Brain Lipids, Lipid Rafts, and Lipid Droplets in Aging and Alzheimer’s Disease. Biomolecules 14, 1362. 10.3390/biom14111362.

46. Islimye, E., Girard, V., and Gould, A.P. (2022). Functions of Stress-Induced Lipid Droplets in the Nervous System. Front Cell Dev Biol 10. 10.3389/fcell.2022.863907.

47. Kao, Y.-C., Ho, P.-C., Tu, Y.-K., Jou, I.-M., and Tsai, K.-J. (2020). Lipids and Alzheimer’s Disease. Int J Mol Sci 21, 1505. 10.3390/ijms21041505.

48. Díaz, M., Fabelo, N., Ferrer, I., and Marín, R. (2018). “Lipid raft aging” in the human frontal cortex during nonpathological aging: gender influences and potential implications in Alzheimer’s disease. Neurobiol Aging 67, 42–52. 10.1016/j.neurobiolaging.2018.02.022.

49. Cutler, R.G., Kelly, J., Storie, K., Pedersen, W.A., Tammara, A., Hatanpaa, K., Troncoso, J.C., and Mattson, M.P. (2004). Involvement of oxidative stress-induced abnormalities in ceramide and cholesterol metabolism in brain aging and Alzheimer’s disease. Proceedings of the National Academy of Sciences 101, 2070–2075. 10.1073/pnas.0305799101.

50. Emre, C., Do, K. V., Jun, B., Hjorth, E., Alcalde, S.G., Kautzmann, M.-A.I., Gordon, W.C., Nilsson, P., Bazan, N.G., and Schultzberg, M. (2021). Age-related changes in brain phospholipids and bioactive lipids in the APP knock-in mouse model of Alzheimer’s disease. Acta Neuropathol Commun 9, 116. 10.1186/s40478-021-01216-4.

51. Tian, G., Kong, Q., Lai, L., Ray-Chaudhury, A., and Lin, C.G. (2010). Increased expression of cholesterol 24S-hydroxylase results in disruption of glial glutamate transporter EAAT2 association with lipid rafts: a potential role in Alzheimer’s disease. J Neurochem 113, 978–989. 10.1111/j.1471-4159.2010.06661.x.

52. Ayciriex, S., Djelti, F., Alves, S., Regazzetti, A., Gaudin, M., Varin, J., Langui, D., Bièche, I., Hudry, E., Dargère, D., et al. (2017). Neuronal Cholesterol Accumulation Induced by Cyp46a1 Down-Regulation in Mouse Hippocampus Disrupts Brain Lipid Homeostasis. Front Mol Neurosci 10. 10.3389/fnmol.2017.00211.

53. Djelti, F., Braudeau, J., Hudry, E., Dhenain, M., Varin, J., Bièche, I., Marquer, C., Chali, F., Ayciriex, S., Auzeil, N., et al. (2015). CYP46A1 inhibition, brain cholesterol accumulation and neurodegeneration pave the way for Alzheimer’s disease. Brain 138, 2383–2398. 10.1093/brain/awv166.

54. Rushworth, J. V., and Hooper, N.M. (2011). Lipid Rafts: Linking Alzheimer′s Amyloid- β Production, Aggregation, and Toxicity at Neuronal Membranes. Int J Alzheimers Dis 2011. 10.4061/2011/603052.

55. Kim, S., Coukos, R., Gao, F., and Krainc, D. (2022). Dysregulation of organelle membrane contact sites in neurological diseases. Neuron 110, 2386–2408. 10.1016/j.neuron.2022.04.020.

56. Vrijsen, S., Vrancx, C., Del Vecchio, M., Swinnen, J. V., Agostinis, P., Winderickx, J., Vangheluwe, P., and Annaert, W. (2022). Inter-organellar Communication in Parkinson’s and Alzheimer’s Disease: Looking Beyond Endoplasmic Reticulum-Mitochondria Contact Sites. Front Neurosci 16. 10.3389/fnins.2022.900338.

57. Goodman, L.D., and Bellen, H.J. (2022). Recent insights into the role of glia and oxidative stress in Alzheimer’s disease gained from Drosophila. Curr Opin Neurobiol 72, 32–38. 10.1016/j.conb.2021.07.012.

58. Liu, L., MacKenzie, K.R., Putluri, N., Maletić-Savatić, M., and Bellen, H.J. (2017). The Glia-Neuron Lactate Shuttle and Elevated ROS Promote Lipid Synthesis in Neurons and Lipid Droplet Accumulation in Glia via APOE/D. Cell Metab 26, 719–737.e6. 10.1016/j.cmet.2017.08.024.

59. Kadyrov, M., Whiley, L., Brown, B., Erickson, K.I., and Holmes, E. (2022). Associations of the Lipidome with Ageing, Cognitive Decline and Exercise Behaviours. Metabolites 12, 822. 10.3390/metabo12090822.

60. Marschallinger, J., Iram, T., Zardeneta, M., Lee, S.E., Lehallier, B., Haney, M.S., Pluvinage, J. V., Mathur, V., Hahn, O., Morgens, D.W., et al. (2020). Lipid-droplet-accumulating microglia represent a dysfunctional and proinflammatory state in the aging brain. Nat Neurosci 23, 194–208. 10.1038/s41593-019-0566-1.

61. Barnett, A.M., Dawkins, L., Zou, J., Mcnair, E., Nikolova, V.D., Moy, S.S., Sutherland, G.T., Stevens, J., Colie, M., Katemboh, K., et al. (2024). Loss of neuronal lysosomal acid lipase drives amyloid pathology in Alzheimer’s disease. 10.1101/2024.06.09.596693.

62. Haney, M.S., Pálovics, R., Munson, C.N., Long, C., Johansson, P.K., Yip, O., Dong, W., Rawat, E., West, E., Schlachetzki, J.C.M., et al. (2024). APOE4/4 is linked to damaging lipid droplets in Alzheimer’s disease microglia. Nature 628, 154–161. 10.1038/s41586-024-07185-7.

63. Ioannou, M.S., Jackson, J., Sheu, S.-H., Chang, C.-L., Weigel, A. V., Liu, H., Pasolli, H.A., Xu, C.S., Pang, S., Matthies, D., et al. (2019). Neuron-Astrocyte Metabolic Coupling Protects against Activity-Induced Fatty Acid Toxicity. Cell 177, 1522–1535.e14. 10.1016/j.cell.2019.04.001.

64. Mi, Y., Qi, G., Vitali, F., Shang, Y., Raikes, A.C., Wang, T., Jin, Y., Brinton, R.D., Gu, H., and Yin, F. (2023). Loss of fatty acid degradation by astrocytic mitochondria triggers neuroinflammation and neurodegeneration. Nat Metab 5, 445–465. 10.1038/s42255-023-00756-4.

65. Qi, G., Mi, Y., Shi, X., Gu, H., Brinton, R.D., and Yin, F. (2021). ApoE4 Impairs Neuron-Astrocyte Coupling of Fatty Acid Metabolism. Cell Rep 34, 108572. 10.1016/j.celrep.2020.108572.

66. Di Paolo, G., and Kim, T.-W. (2011). Linking lipids to Alzheimer’s disease: cholesterol and beyond. Nat Rev Neurosci 12, 284–296. 10.1038/nrn3012.

67. Fu, Y., He, Y., Phan, K., Pickford, R., Kim, Y.-B., Dzamko, N., Halliday, G.M., and Kim, W.S. (2022). Sex-specific lipid dysregulation in the *Abca7* knockout mouse brain. Brain Commun 4. 10.1093/braincomms/fcac120.

68. Dib, S., Pahnke, J., and Gosselet, F. (2021). Role of ABCA7 in Human Health and in Alzheimer’s Disease. Int J Mol Sci 22, 4603. 10.3390/ijms22094603.

69. Dong, Y., Harlan, B.A., and Brewer, G.J. (2021). When aging switches on Alzheimer’s. Aging 13, 13376–13377. 10.18632/aging.203085.

70. Geltinger, F., Schartel, L., Wiederstein, M., Tevini, J., Aigner, E., Felder, T.K., and Rinnerthaler, M. (2020). Friend or Foe: Lipid Droplets as Organelles for Protein and Lipid Storage in Cellular Stress Response, Aging and Disease. Molecules 25, 5053. 10.3390/molecules25215053.

71. Smolič, T., Tavčar, P., Horvat, A., Černe, U., Halužan Vasle, A., Tratnjek, L., Kreft, M.E., Scholz, N., Matis, M., Petan, T., et al. (2021). Astrocytes in stress accumulate lipid droplets. Glia 69, 1540–1562. 10.1002/glia.23978.

72. Brewer, G.J., and Torricelli, J.R. (2007). Isolation and culture of adult neurons and neurospheres. Nat Protoc 2, 1490–1498. 10.1038/nprot.2007.207.

