## Supplementary figures and images for "AD-genes and Aging Increase Count and Size of Lipid Droplets, Accompanied by Accumulation of Neutral Lipids Across Compartments in Hippocampal Neurons"

### Supplementary Figure 1

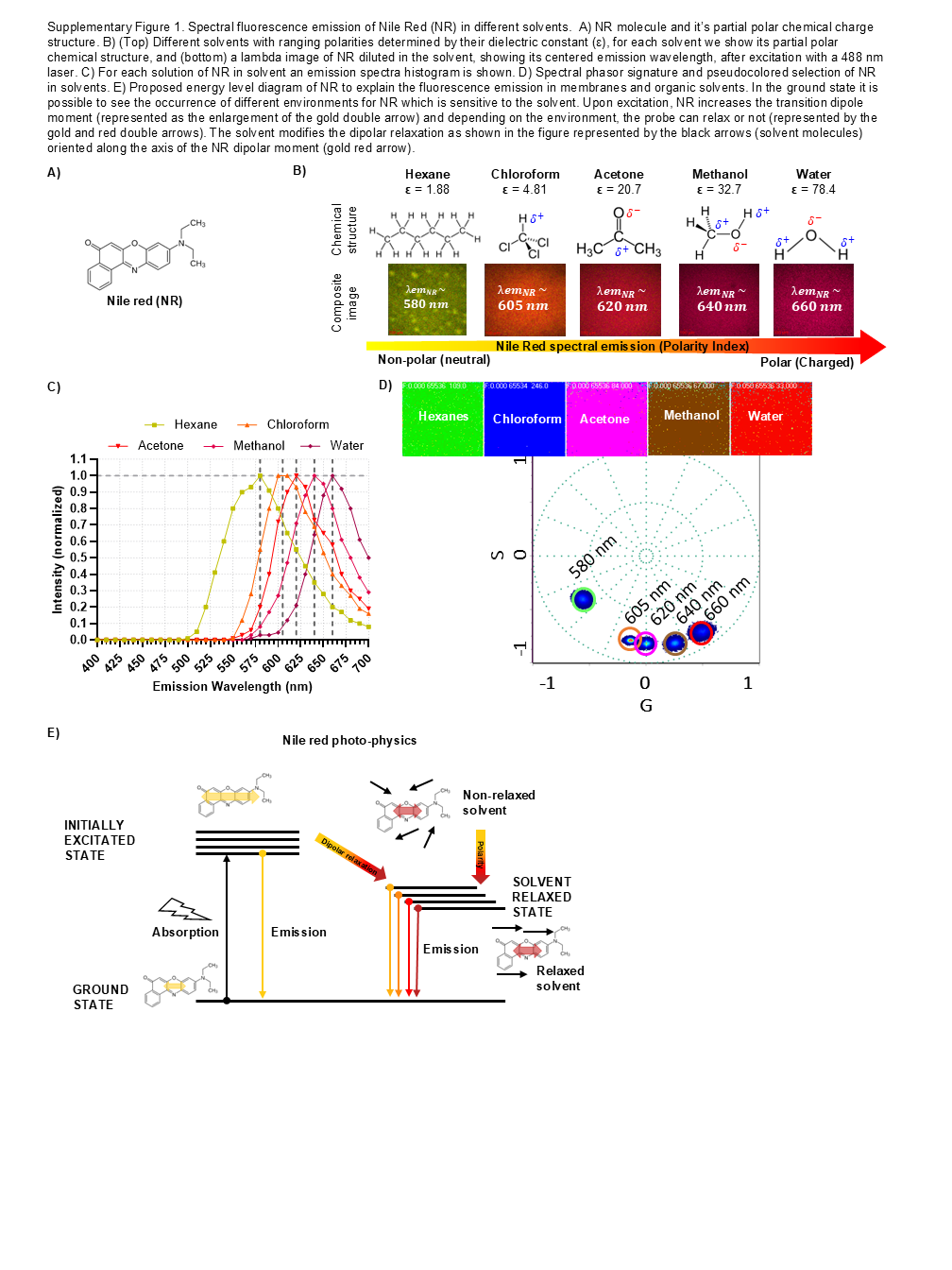

### Supplementary Figure 2

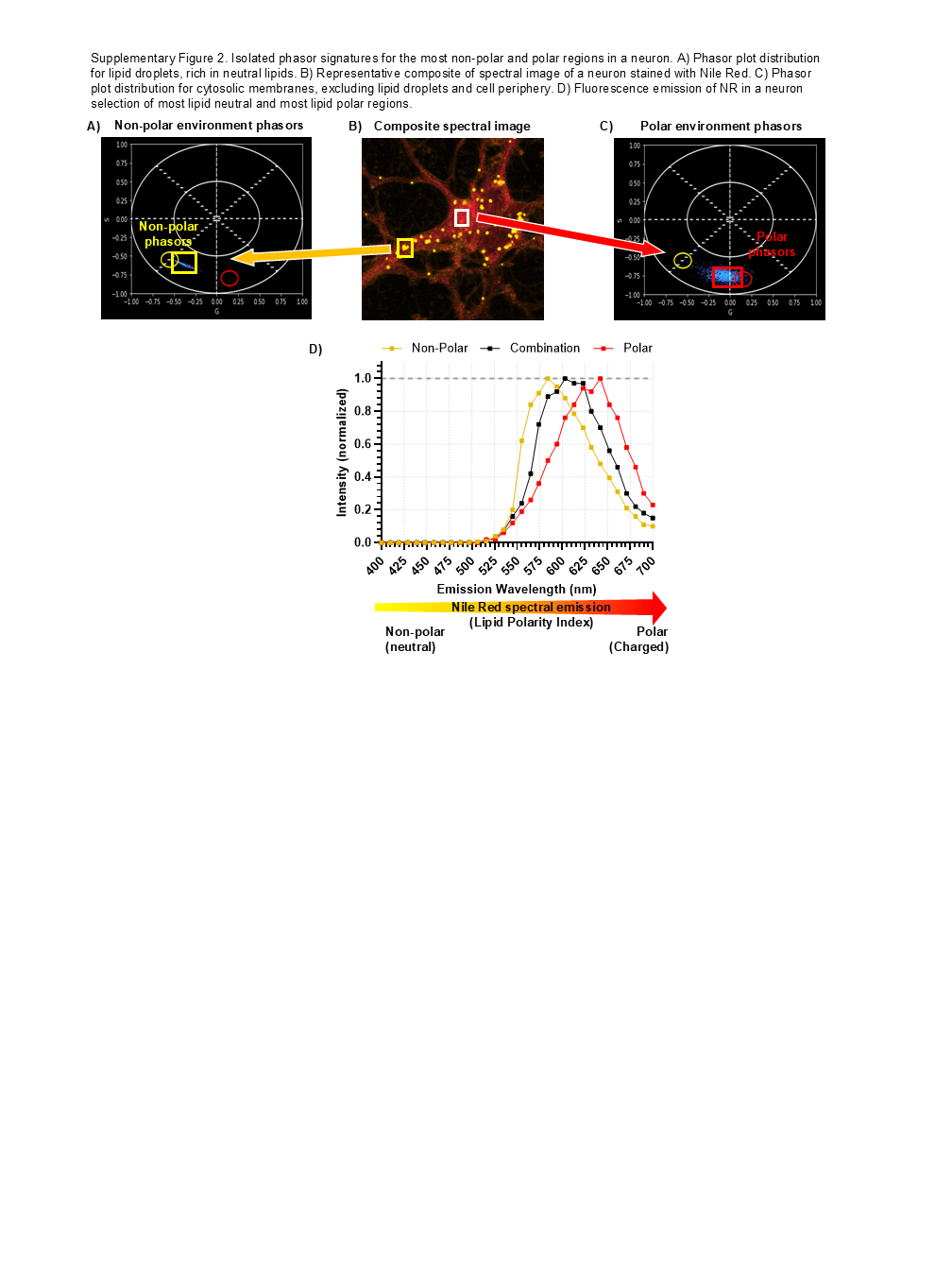

### Supplementary Figure 3

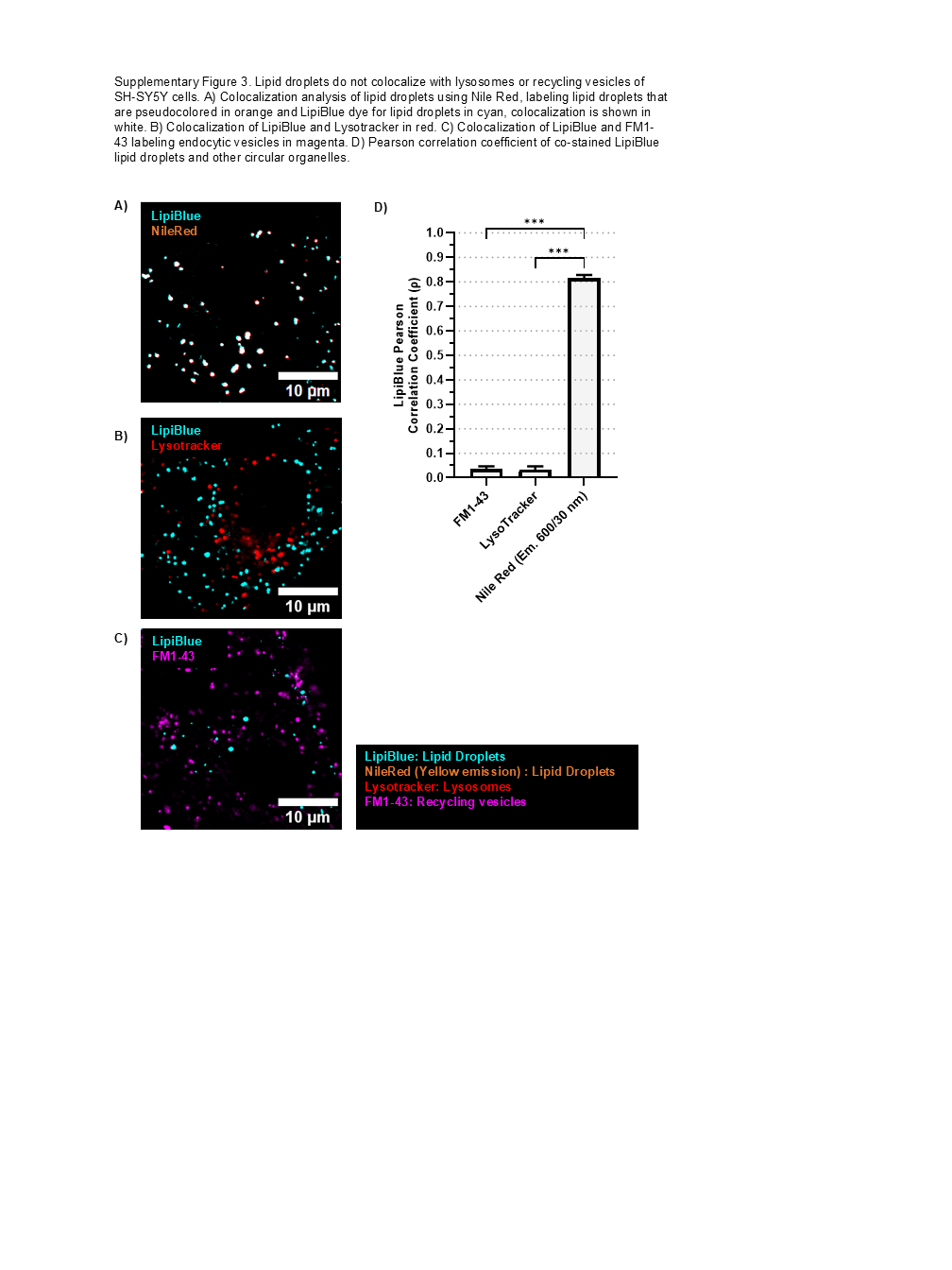

### Supplementary Figure 4

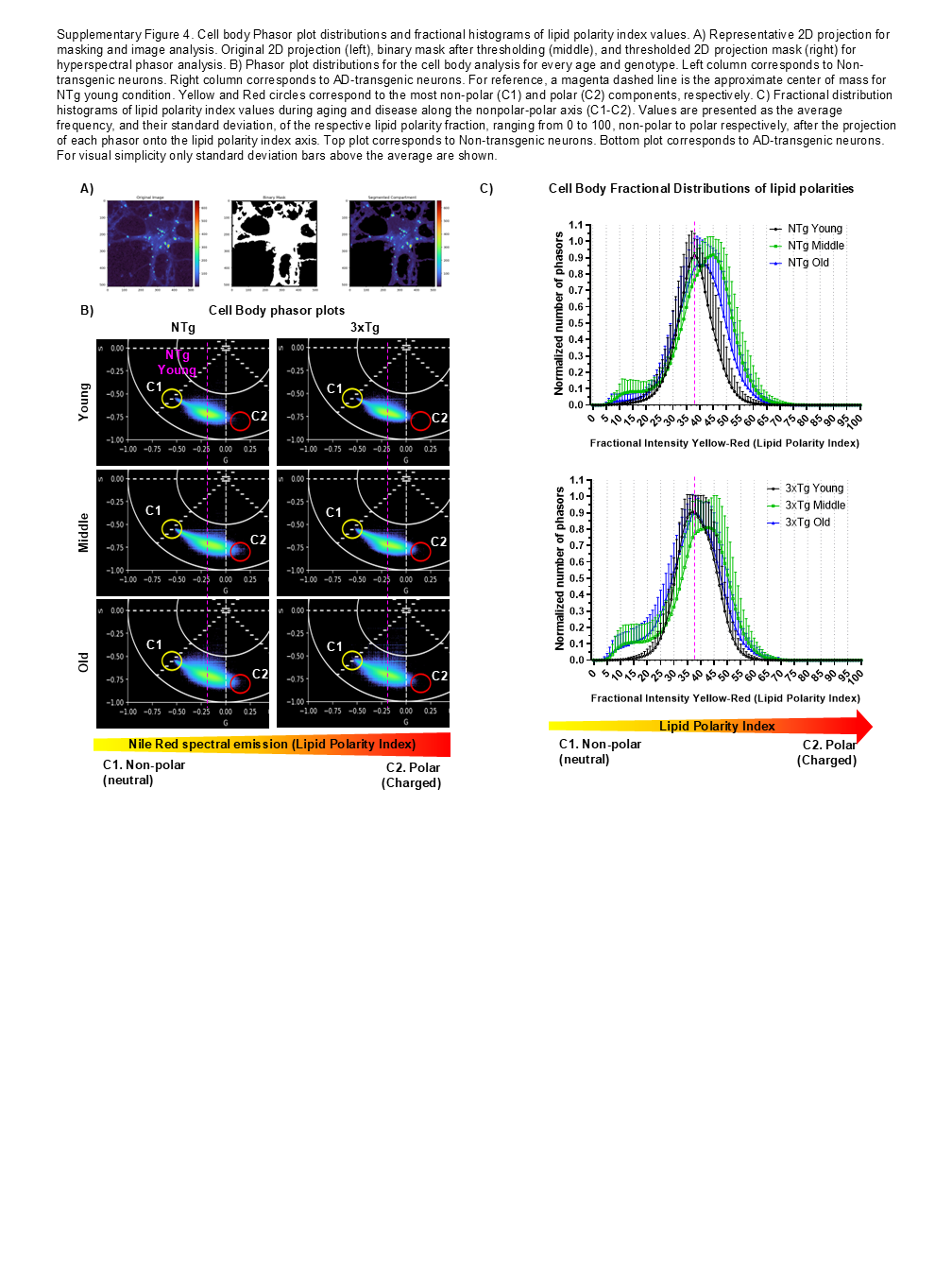

### Supplementary Figure 5

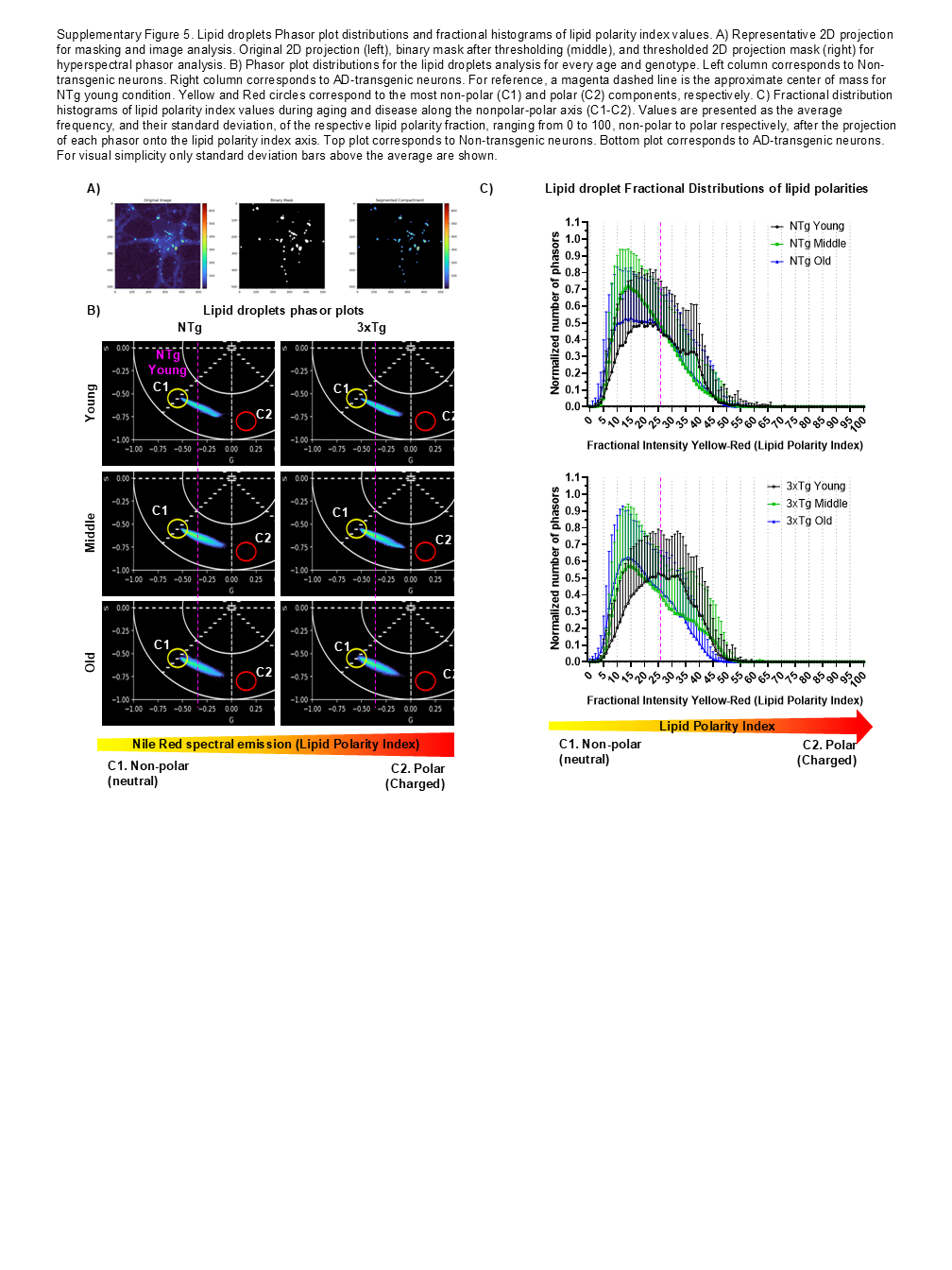

### Supplementary Figure 6

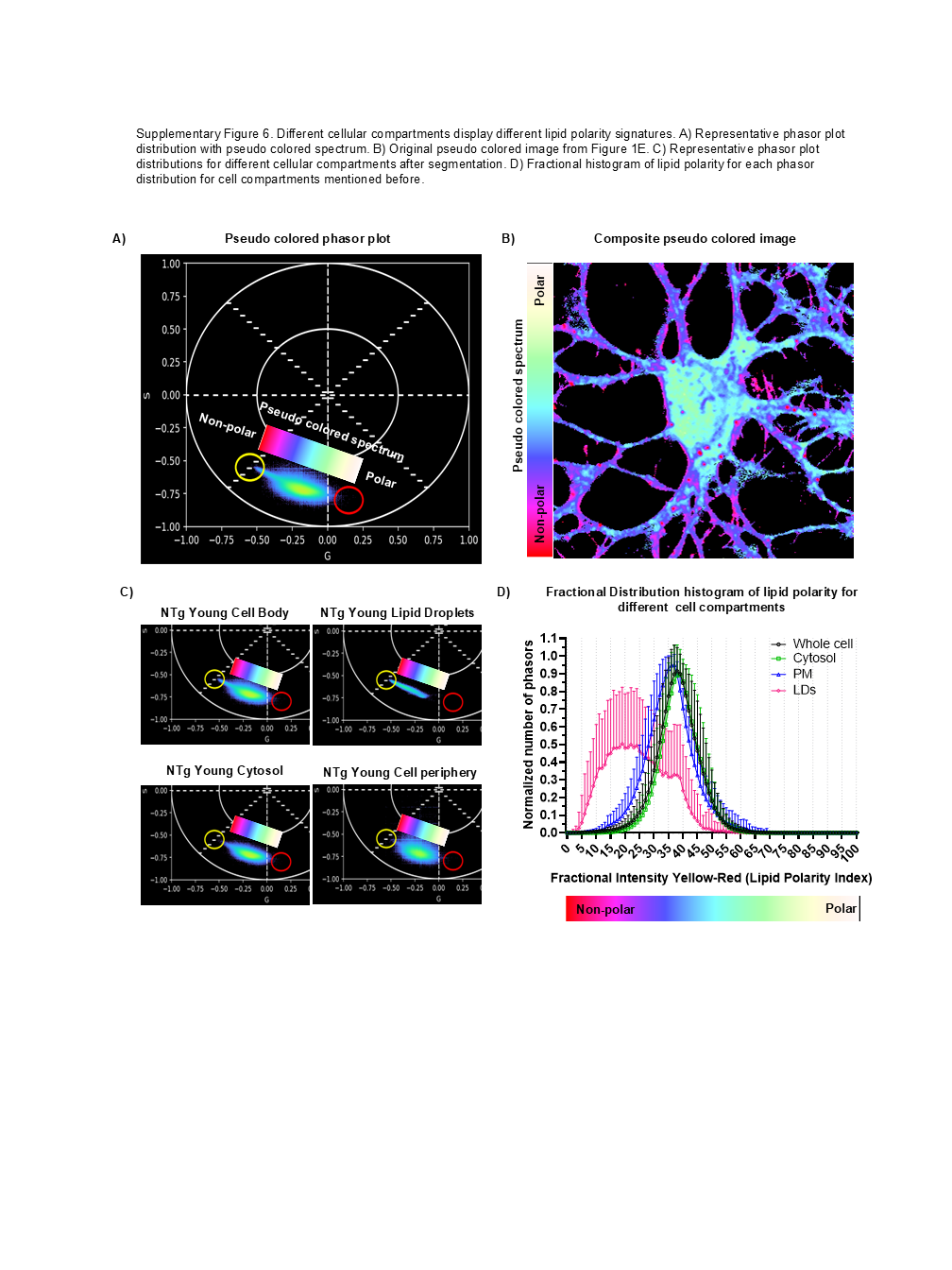

### Supplementary Figure 7

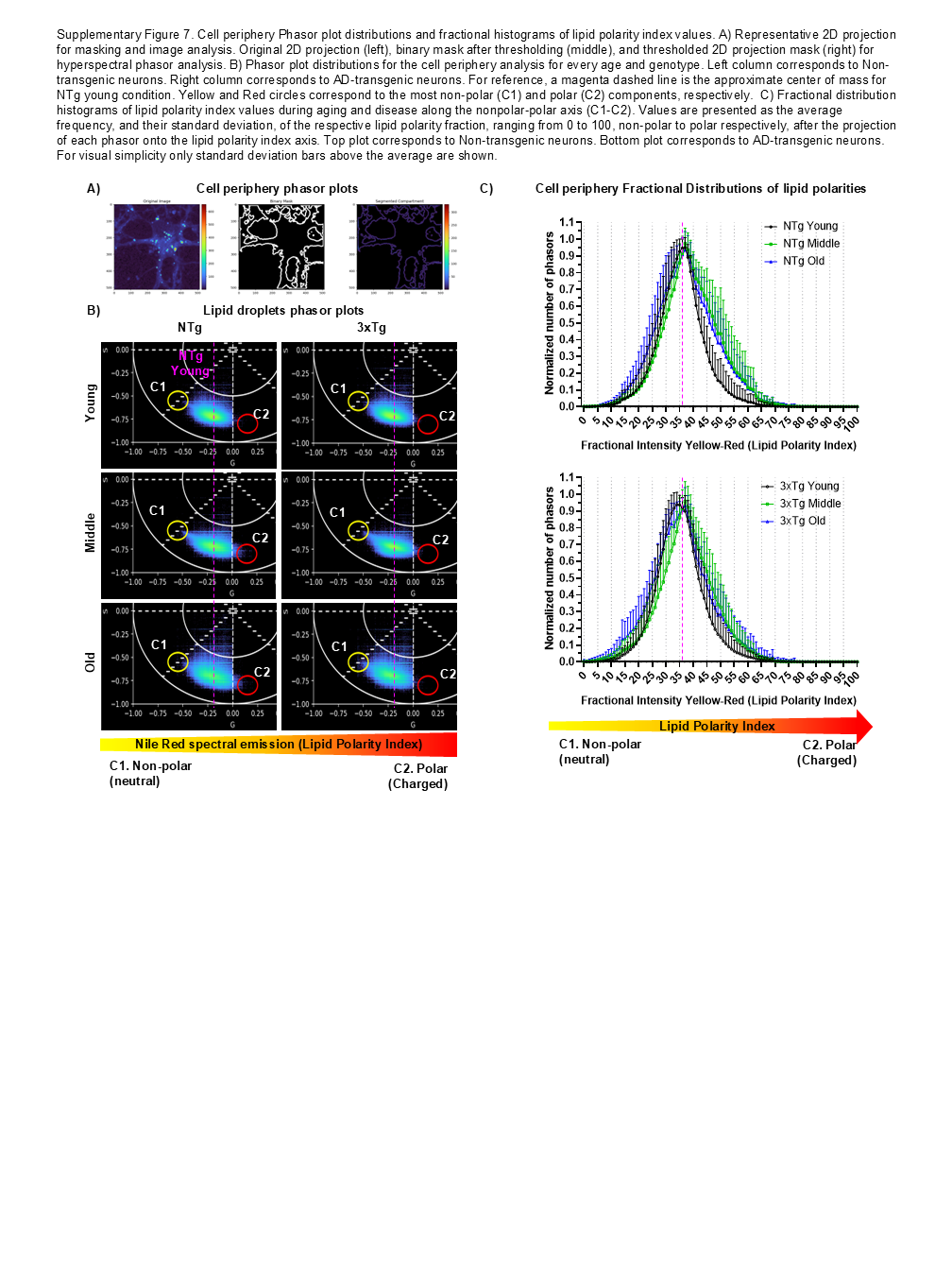

### Supplementary Figure 8

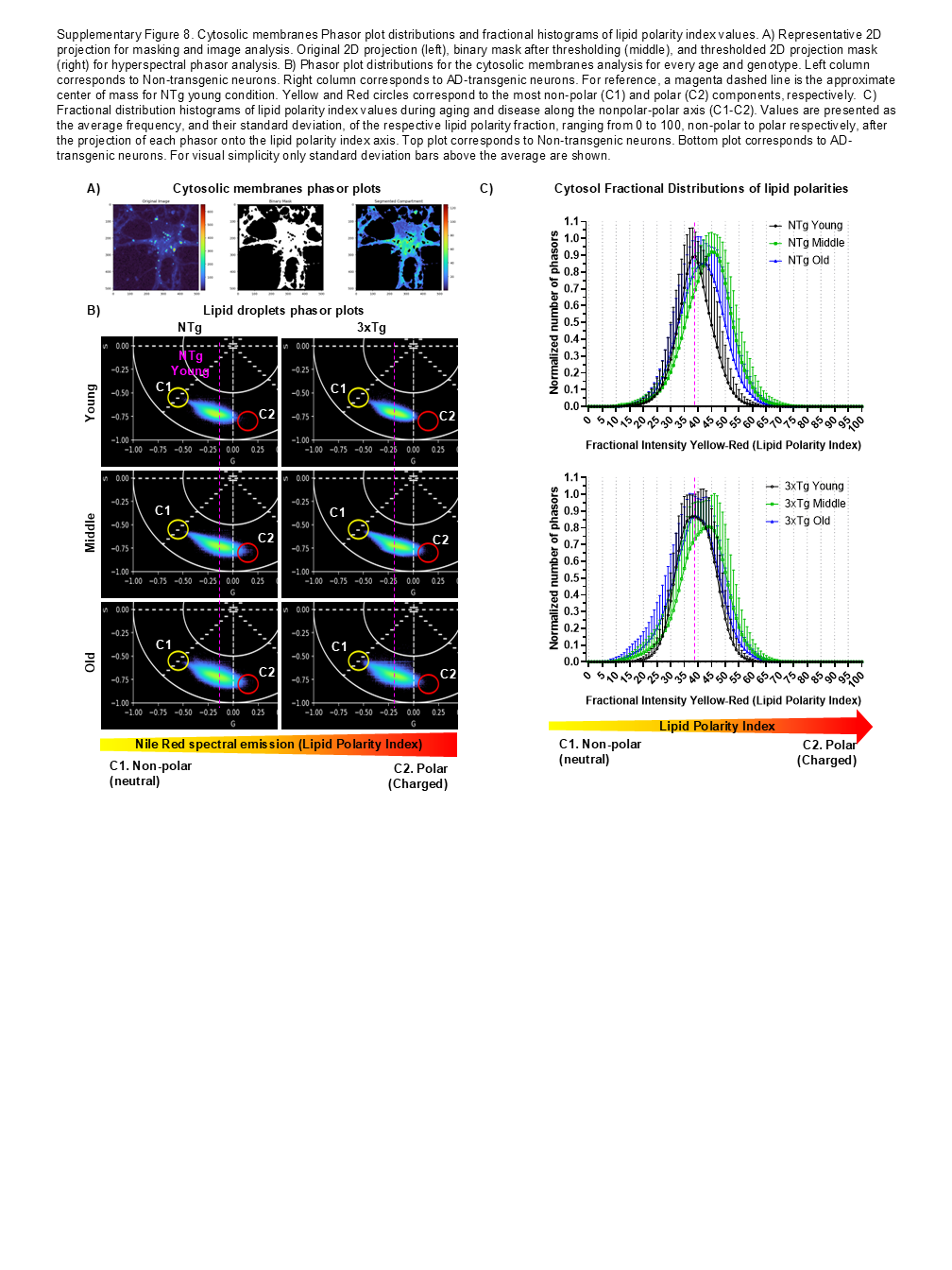

### Supplementary Figure 9

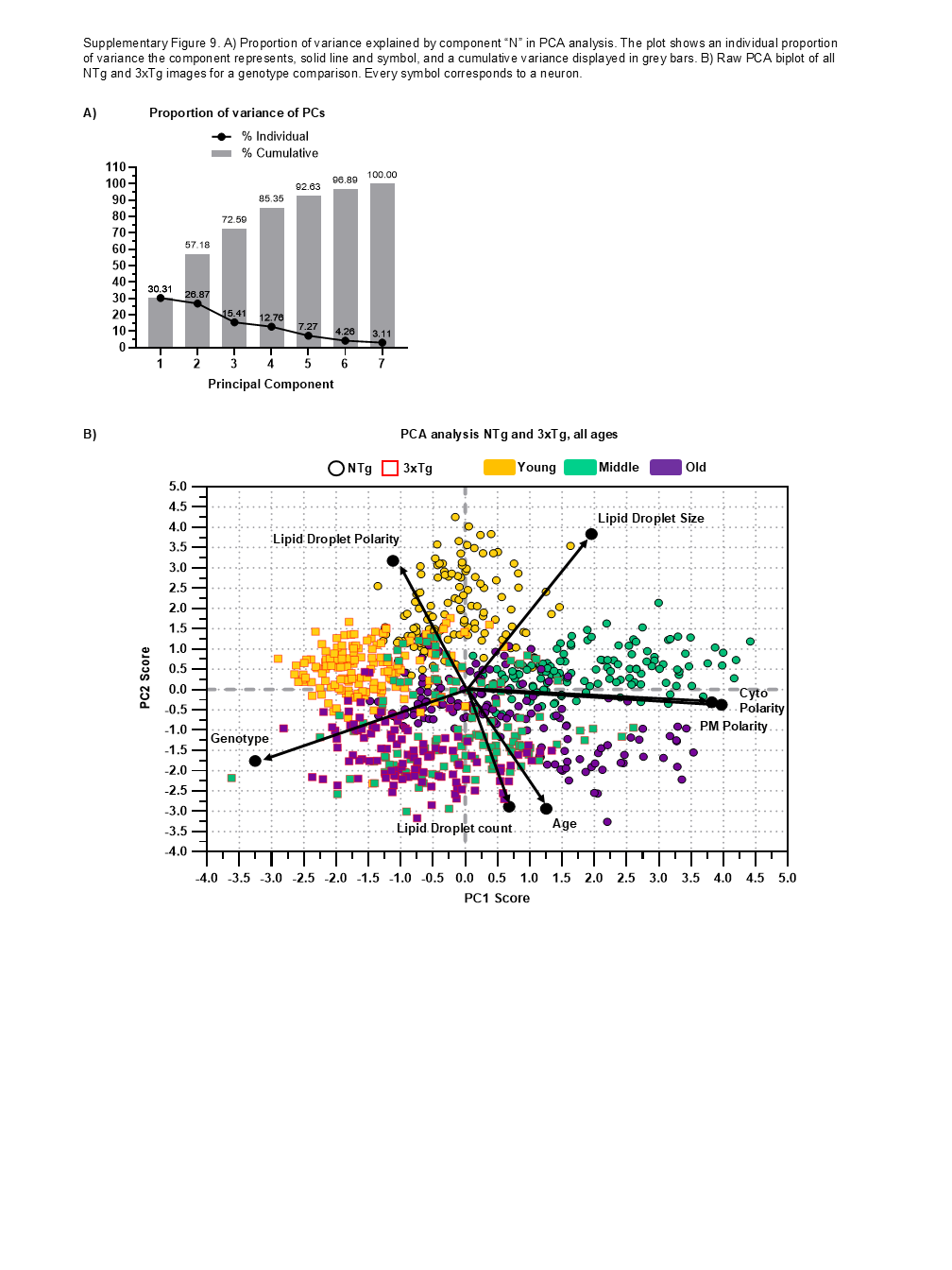
