## Supplementary Table 1 for "AD-genes and Aging Increase Count and Size of Lipid Droplets, Accompanied by Accumulation of Neutral Lipids Across Compartments in Hippocampal Neurons"

**Supplementary Table1.** Loading Table in Principal Component Analysis. Principal Components identify key cellular patterns of lipid changes that distinguish young from old and AD-genotypes. A global PCA reveals key sources of variance in our data. The following variables were used: genotype, age, cytosolic polarity, cell periphery (PM) polarity, LD polarity, LD average count, and LD average size. PCA decomposed their variance into seven principal components. Based on Parallel Analysis, PC1 (30%), PC2 (26%), and PC3 (15%) were selected, together capturing 72% of the total variance (Supplementary Figure 9A). A loadings table from the PCA analysis describes the correlation between each original variable and each principal component (PC). A larger loading (positive or negative magnitude) indicates a stronger relationship between the variable and the PC. The sign of the loading indicates whether the variable and the PC are positively or negatively correlated. A positive loading indicates the contribution of a variable to the principal component, whereas a negative loading indicates the absence of a contribution of a variable to the principal component.

| Variable | PC1 | PC2 | PC3 |
|---|---|---|---|
| LD LP | -0.20 | **0.57** | **-0.60** |
| LD count | 0.14 | **-0.65** | -0.11 |
| LD size | **0.42** | **0.69** | 0.20 |
| Cyto LP | **0.83** | -0.07 | **-0.40** |
| PM LP | **0.86** | -0.08 | **-0.30** |
| Genotype | **-0.58** | **-0.40** | **-0.58** |
| Age | **0.27** | **-0.66** | **0.24** |
